# Single-cell analysis resolves the genetic program of primary vascular system formation in hybrid poplar

**DOI:** 10.1101/2021.12.28.474348

**Authors:** Daniel Conde, Paolo M. Triozzi, Wendell J. Pereira, Henry W. Schmidt, Kelly M. Balmant, Sara A. Knaack, Sushmita Roy, Christopher Dervinis, Matias Kirst

**Affiliations:** School of Forest, Fisheries and Geomatics Sciences, University of Florida, Gainesville, FL 32611, USA; Wisconsin Institute for Discovery, University of Wisconsin, Madison, Wisconsin 53715, WI 53715, USA; Department of Computer Sciences, University of Wisconsin, Madison, WI 53792, USA; Department of Biostatistics and Medical Informatics, University of Wisconsin, Madison, WI 53792, USA; Genetics Institute, University of Florida, Gainesville, FL 32611, USA

## Abstract

Despite the enormous potential of novel approaches to explore gene expression at a single-cell level, we lack a high-resolution and cell type-specific gene expression map of the shoot apex in woody perennials. We use single-nuclei RNA sequencing to determine the cell type-specific transcriptome of the *Populus* vegetative shoot apex. We identified highly heterogeneous cell populations clustered into seven broad groups represented by 18 transcriptionally distinct cell clusters. Next, we established the developmental trajectories of epidermal cells, leaf mesophyll, and vascular tissue. Motivated by the high similarities between *Populus* and *Arabidopsis* cell population in the vegetative apex, we created and applied a pipeline for interspecific single-cell expression data integration. We contrasted the developmental trajectories of primary phloem and xylem formation in both species, establishing the first comparison of primary vascular development between a model annual herbaceous and a woody perennial plant species. Our results offer a valuable resource for investigating the basic principles underlying cell division and differentiation conserved between herbaceous and perennial species, which also allows the evaluation of the divergencies at single-cell resolution.

**One Sentence Summary:** Single-cell RNA-seq of the vegetative shoot apex resolved the genetic program of primary vascular development in hybrid poplar.

## INTRODUCTION

A multicellular organism’s capacity to develop body structures is dependent on pluripotent stem cells. In plants, these cells are present in meristematic tissues, including the shoot apical meristem (SAM). Except for the hypocotyl and cotyledons, the SAM is responsible for generating all aerial parts of the plant. The meristem must balance the self-renewal of a reservoir of stem cells and organ initiation to fulfill this function. Most of the knowledge about the signaling networks that regulate this balance came from the model species *Arabidopsis*. The *Arabidopsis* SAM is divided into three cell layers. The L1 layer derivatives give rise to the epidermis of shoots, leaves, and flowers, whereas the L2 layer provides the mesophyll tissue and germ cells, and the L3 layer contributes to the vascular tissues and pith (*1*). The *Arabidopsis* SAM can also be divided into distinct cell domains or zones. Stem cells reside in the central zone (CZ). The peripheral zone (PZ), where cells divide more frequently than in the CZ, is responsible for organ initiation. The rib meristem (RM) gives rise to central tissues of the shoot axis (*2*, *3*).

The stem cells of the vegetative shoot apex undergo divisions to generate transitioning cells that eventually differentiate into lateral organs such as leaves and vasculature. While well explored in the annual herbaceous plant model *Arabidopsis*, we lack an understanding of the origin and trajectory of cell lineages that give rise to shoot apex cells in woody perennial species such as *Populus* sp. Previous profiling studies in *Arabidopsis* used reporter genes to purify domain-specific cell populations (*4*, *5*), an approach that underestimates cell complexity as it restricts the analysis to the expression domains of these markers. The development of microfluidic-based single-cell RNA sequencing (scRNA-seq) methods established the foundation for quantifying the complete transcriptome at cellular resolution and enabled the characterization of cell subpopulations in heterogeneous tissues in plants (*6*–*8*). Moreover, scRNA-seq analysis allows ordering individual cells in a pseudotime trajectory to reveal the lineages that determine plant form. This approach allowed, in *Arabidopsis*, the discovery of the developmental trajectories from SAM proliferating cells to the formation of new organs (*1*). This approach has also been used in *Arabidopsis* to explore root development (*9*–*12*). However, methods of protoplast isolation used in *Arabidopsis* are not immediately applicable to many tissues and species. Consequently, a high-resolution and cell type-specific gene expression map of the shoot apex of woody perennials is still lacking, resulting in a limited understanding of the regulatory mechanism involved in the organ differentiation from SAM stem cells.

Here we applied a novel approach to isolate nuclei from complex plant tissues (*13*), to dissect the transcriptome profile of the hybrid poplar (*Populus tremula* × *alba*) vegetative shoot apex at single-cell resolution. We inferred the developmental trajectories that occur during the generation of new tissues in the SAM. We then assessed the development of the primary vasculature, a process largely unexplored in woody perennial species, and identified regulators of the vascular tissue differentiation. Finally, we created and applied a pipeline for the interspecific comparison of single-cell transcriptome data. We contrasted the developmental trajectories of primary phloem and xylem formation in *Arabidopsis* and *Populus* with this pipeline, establishing the first comparison between primary vasculature development at the single-cell level between a model annual herbaceous and a woody perennial plant species.

## RESULTS

### An atlas of *Populus* vegetative shoot apex cells

We harvested 20 shoot apices from 3-weeks-old *in vitro* grown hybrid poplar (*Populus tremula* × *alba*) plants and performed high-throughput, microfluidic-based single-nuclei RNA sequencing (snRNA-seq) using the 10× Genomics Chromium technology as we described previously (*13*). For all quality control and analytical steps, we used Asc-Seurat (*14*), a comprehensive web application that encapsulates a series of tools for scRNA-seq data analysis. We removed potentially empty GEMs (containing only ambient RNA) by requiring a minimum of 1,000 unique molecular identifiers (UMI)-tagged transcripts detected per nucleus and excluded those nuclei with more than 7,000 (Table S1, Fig. S1). We captured 8,324 high-quality nuclei. An average of 3,618 unique molecular identifiers (UMIs), corresponding to the expression of 2,477 genes, were detected per nucleus. Overall, 31,214 genes were detected (Table S1). Principal component analysis and unsupervised expression data analyses uncovered 18 distinct clusters (Fig. 1A, Fig. S1). To annotate each cluster, we assessed the abundance of transcripts of poplar homologs to *Arabidopsis* well-known cell type marker genes for the different domains of the vegetative or reproductive shoot apex (*1*, *4*, *5*) (Table S2). We also evaluated the expression of homologous genes that, in *Arabidopsis*, are mainly expressed at different cell types of the vegetative and the reproductive shoot apex (*4*, *5*). Moreover, we identified all the cell markers for each cluster detected in the poplar shoot apex (Table S3) and explored those whose biological functions or expression patterns have previously been well characterized. Based on these three sources of information, we determined the poplar tissue-specific markers used to annotate the clusters (Table S2). Clustering annotation revealed populations of seven cell types: leaf primordia (LF), mesophyll cells (MC), epidermal cells (EC), shoot meristematic cells (SMC), proliferating cells (PC), vascular cells (VC), and companion cells (CC) (Fig. 1 A-D).

**Fig. 1.**
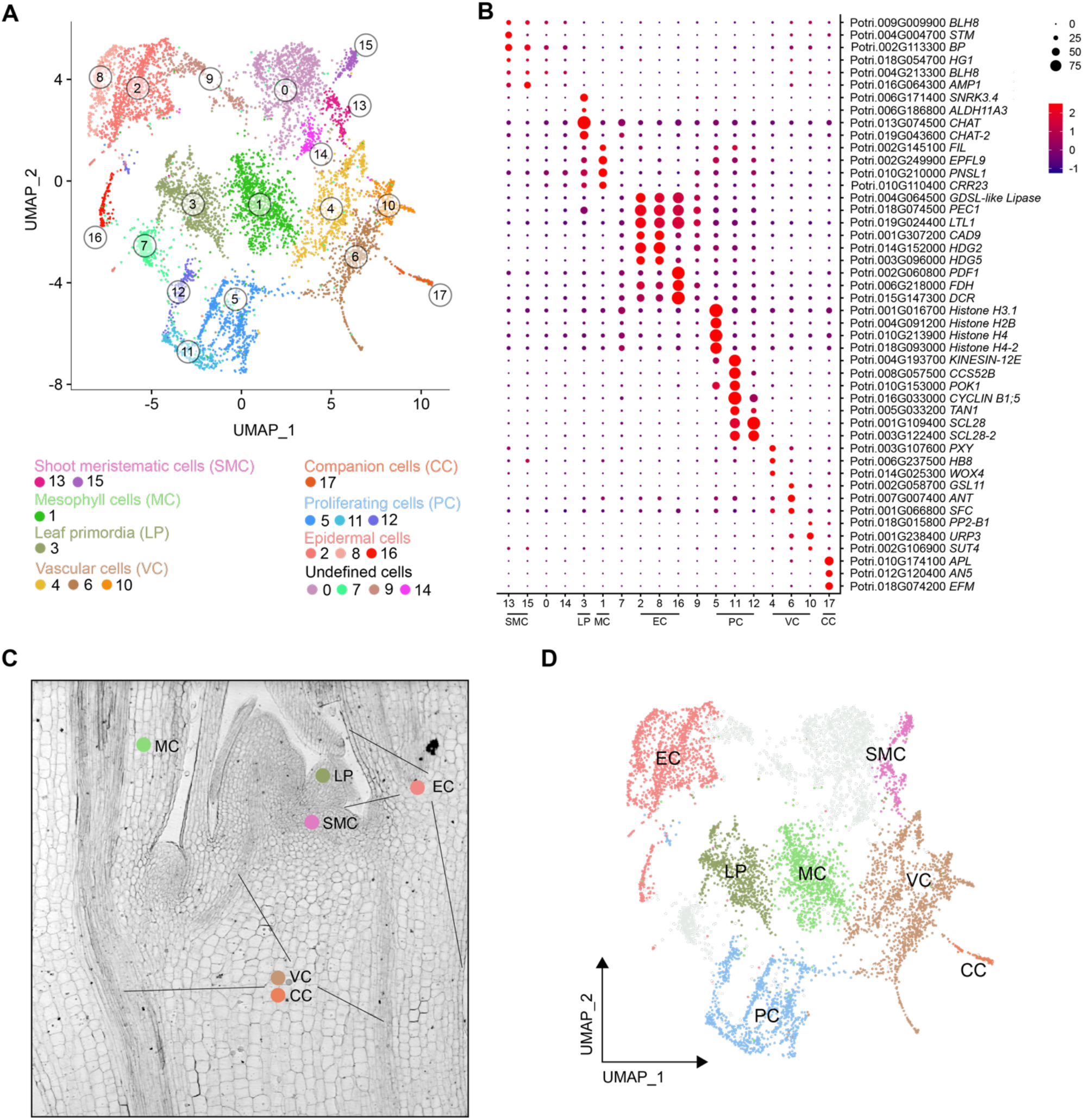
Characterization of cell populations in the poplar vegetative shoot apex. **(A)** Visualization of the 18 cell clusters using UMAP. Dots, individual cells; n = 8,324 cells; color, cell clusters. **(B)** Expression pattern of previously characterized cell-type marker genes in the poplar apex cell clusters. Dot diameter, the proportion of cluster cells expressing a given gene. The complete list of genes used for cluster annotation is given in table S2. **(C)** Longitudinal section of a poplar apex, showing the spatial location of the population of annotated cells. **(D)** Visualization of 7 cell types using UMAP. Color, population types.

The MC population consisted of one cluster (cluster 1) (Fig. 1A), in which poplar homologs to *Arabidopsis* markers for the mesophyll, such as *EPIDERMAL PATTERNING FACTOR LIKE-9* (*EPFL9*) (*15*), *CHLORORESPIRATORY REDUCTION 23* (*CRR23*) (*1*), or *PHOTOSYNTHETIC NDH SUBCOMPLEX L1* (*PNSL1*) (*1*) were predominantly expressed (Fig. 1B, Table S2). Moreover, photosynthesis and small molecule metabolic processes are the most enriched gene ontologies in cluster 1 (Table S4). The epidermal-specific genes *Li-TOLERANT LIPASE 1* (*LTL1*) (*4*), *HOMEODOMAIN GLABROUS 2* (*HDG2*) (*4*), and *PROTODERMAL FACTOR1* (PDF1) (*1*), were detected in the EC population (i.e., L1 layer), which consisted of clusters 2, 8, and 16 (Fig. 1B). The SMC population consisted of two clusters, 13 and 15 (Fig. 1B). Transcripts for poplar homologs to the *Arabidopsis SHOOT MERISTEMLESS* (*STM*) and *BREVIPEDICELLUS* (*BP*), which are required for the establishment and maintenance of the *Arabidopsis* SAM (*1*, *16*), and other related homeodomain genes, *HOMEOBOX GENE 1* (*HG1*) (*17*) and *BEL1-LIKE HOMEODOMAIN 8* (*BLH8*) (*18*), were highly enriched in this cell populations (Fig. 1B). We annotated three clusters (5, 11, 12) as the PC population because transcripts for homologs to cell-cycle-related genes such as *HISTONE H4* (*H4*), *CELL CYCLE SWITCH PROTEIN 52 B* (*CCS52B*), *CYCLIN B1;5* (*CYCA1;1*) or *SCARECROW-LIKE 28* (*SCL28*) (*19*, *20*) were overrepresented (Fig. 1B). The VC population was composed of four clusters (4, 6, 10, and 17) (Fig. 1A), in which genes involved in xylem and phloem differentiation were expressed. For instance, transcripts of the poplar homolog to the xylem gene *PHLOEM INTERCALATED WITH XYLEM* (PXY) (*21*) and the phloem gene *PHLOEM PROTEIN 2-B1* (*PP2-B1*) (*22*) were markedly overrepresented in clusters 4 and 10, respectively (Fig. 1B). Homologs to markers for cambium stem cells, such as *AINTEGUMENTA* (*ANT*) (*21*), were enriched in cluster 6. The CC population (cluster 17) was observed in a highly distinct cluster on the UMAP plot, agreeing with their specific physiological functionalities and unique expression profiles (Fig. 1A). Transcripts for poplar homologs to CC marker genes such as *ARATHNICTABA 5* (*AN5*) and *EARLY FLOWERING MYB* (*EFM*) (*1*) are highly accumulated in cluster 17 (Fig. 1B). Finally, we annotated cluster 3 as leaf primordia, based on the enrichment in genes expressed in the domains of *KANADI 1* (*KAN1*) and *FILAMENTOUS FLOWER* (*FIL*) in the *Arabidopsis* vegetative and reproductive SAM (*4*, *5*) (Fig. 1B, Table S2). The above results indicate that the vegetative shoot apex is composed of highly heterogeneous cells and that many regulatory mechanisms of the vegetative SAM and shoot apex are conserved in *Populus* and *Arabidopsis*.

### Developmental trajectories of the epidermis, mesophyll, and vascular tissue

Single-cell RNA sequencing has recently allowed the identification of the developmental trajectories of the epidermis, mesophyll, and vasculature differentiation in the *Arabidopsis* vegetative shoot apex (*1*). As previously described for the *Arabidopsis* shoot apex, we identified a large population of proliferating cells in *Populus*. Following the same approach performed in (*1*), we dissected the PC population to identify the specific sub-groups of EC (L1 layer), MC (L2 layer), and VC (L3 layer) proliferating cells. For this, clusters 5, 11, and 12 (Fig. 1A) were re-grouped at higher resolution, revealing seven sub-clusters (Fig. 2A). Sub-clusters 0 and 5 exhibited high transcript levels of *EPLF9* (a marker of MC) and *CHAT* (a marker of LP), respectively (Fig. 2B). Sub-clusters 2 and 6 showed high expression levels of the epidermal cell markers *PDF1*, *HDG2*, or *FIDDLEHEAD* (*FDH*) (*1*) (Fig. 2B). In contrast, the vascular meristem markers *LIKE AUXIN RESISTANT 2* (*LAX2*) (*23*) and *HOMEOBOX GENE 8* (*HB8*) (*24*) were highly expressed in sub-cluster 4 (Fig. 2B). The transcript profile of these marker genes indicates that the PC population contains the cells that differentiate into the epidermis, mesophyll, and vascular cells. Finally, we delineated the developmental trajectories that give rise to these cells. We re-grouped the EC (clusters 2, 8, and 16; Fig. 1A), MC (cluster 1; Fig. 1A), and VC clusters (clusters 4, 6, 10, and 16; Fig. 1A) and included the corresponding proliferating cell sub-clusters (Fig. 2A) of EC, MC, and VC respectively (Fig. 2C). We applied Slingshot (*25*) to infer the developmental trajectory for the epidermis, mesophyll, and vascular tissue differentiation (Fig. 2D). Identifying the proliferating cells with transcriptional signatures of EC, MC, and VC populations paved the way to investigate how SAM stem cells differentiate into the distinct new tissues formed in the poplar shoot apex.

**Fig. 2.**
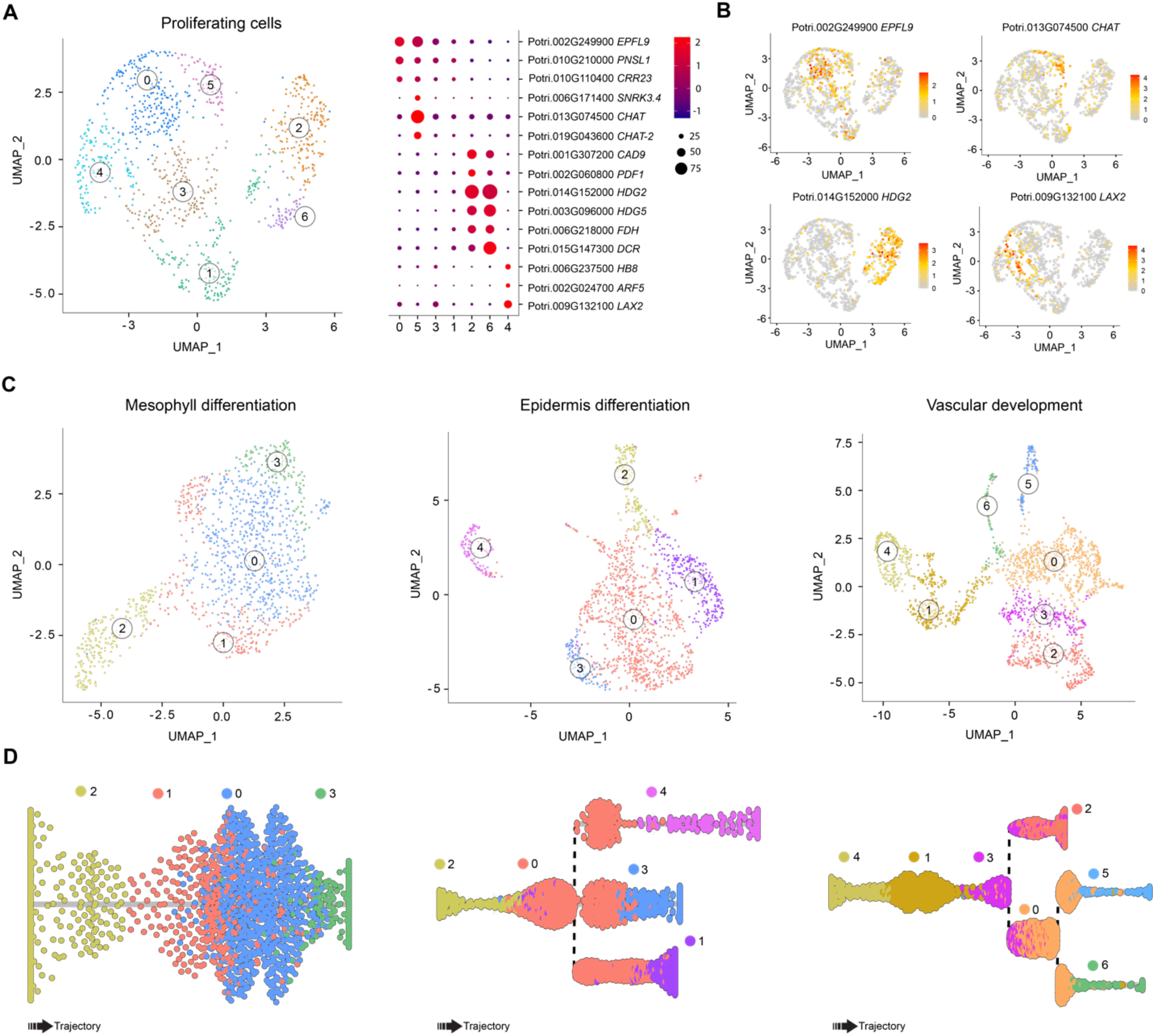
Characterization of the PC population and developmental trajectories for MC, EC, and VC. **(A)** Sub-clustering of clusters 5, 11, and 12 reveals cell-type-specific proliferating cell (PC) populations. **(B)** Expression pattern of well-characterized marker genes revealed the leaf primordia (LP) and mesophyll (MC)-specific (sub-clusters 5 and 0), vascular (VC)-specific (sub-cluster 4), and epidermis (EC)-specific (sub-cluster 2 and 6) cell clusters within the PC population. **(C)** Re-clustering of MC, EC, and VC clusters with their corresponding sub-clusters of PC population, and **(D)** Slingshot analysis showing the position of the sub-clusters during the epidermis, mesophyll, and vascular tissue differentiation trajectory or pseudotime.

### Integration and annotation of *Populus* and *Arabidopsis* vasculature single-cell data

The similarity between cell-type populations identified in *Populus* and *Arabidopsis* shoot apex motivated the exploration of the conservation and divergence of the molecular mechanisms that govern cell differentiation process between these two species, focusing on the primary vascular development. For this comparison, we used the single-cell gene expression data of the shoot apex vasculature of *Arabidopsis* (*1*). Clustering of vascular cells from both species showed a similar cell population structure (Fig. S2A), according to the expression patterns of previously well-characterized markers for proliferating cells, phloem, xylem, and companion cells (Fig. S2B). Integrating both datasets requires a one-to-one homologous gene relationship between *Populus* and *Arabidopsis*. The complex history of whole-genome duplications, chromosomal rearrangements, and tandem duplications in *Populus* resulted in many duplicated genes, which implies that there are several *Populus* homologs for each *Arabidopsis* gene (*26*). To create a robust one-to-one homologous gene list required for the integration, we applied a phylogenomic approach to define a gene set of 9,842 *Arabidopsis* and *Populus* pairs of homologs (Table S5). The expression of these genes was used in Seurat to integrate the single-cell expression data of *Populus* and *Arabidopsis* shoot apex vasculature. Homology between the remaining genes from both species was based on the most recently inferred relationships, available in Phytozome (*27*) (Table S6). After the data integration, we used this complete list to explore conserved and divergent pathways. The UMAP visualization showed that most cells were distributed in common cell-type clusters between both species (Fig. 3A, Fig. S3). Only two clusters enriched in stress responding genes and one small group of cells were specific to *Arabidopsis* (Fig. S3 A, B). Overall, vascular cells were distributed in 12 clusters after the integration. We identified 3 clusters (2, 6, and 7) containing the proliferating cells, based on the expression of cell-cycle related genes such as *CCS52B* or *HISTONE 2B*. Based on the expression pattern of markers used in *Arabidopsis* (*1*) for the sieve elements (*ALTERED PHLOEM DEVELOPMENT* (*APL*) and *SIEVE ELEMENT OCCLUSION-RELATED 1* (*SEOR1*)) and tracheary elements (*VASCULAR RELATED NAC-DOMAIN PROTEIN 1* (*VND1*), *TARGET OF MONOTEROS 5-LIKE 1* (*T5L1*)), we annotated clusters 10 and 5 as phloem sieve elements and xylem tracheary element cells, respectively (Fig. 3B). Next, we used Dynverse to establish the overall developmental trajectory of the integrated vasculature data (Fig. S3C) and determine the clusters associated with cell lineages that result in the formation of sieve and tracheary elements. The overall developmental trajectory pointed to clusters 8 and 10 as being involved in sieve element trajectory, while 8, 0, and 5 participate in tracheary element differentiation (Fig. S3C).

**Fig. 3.**
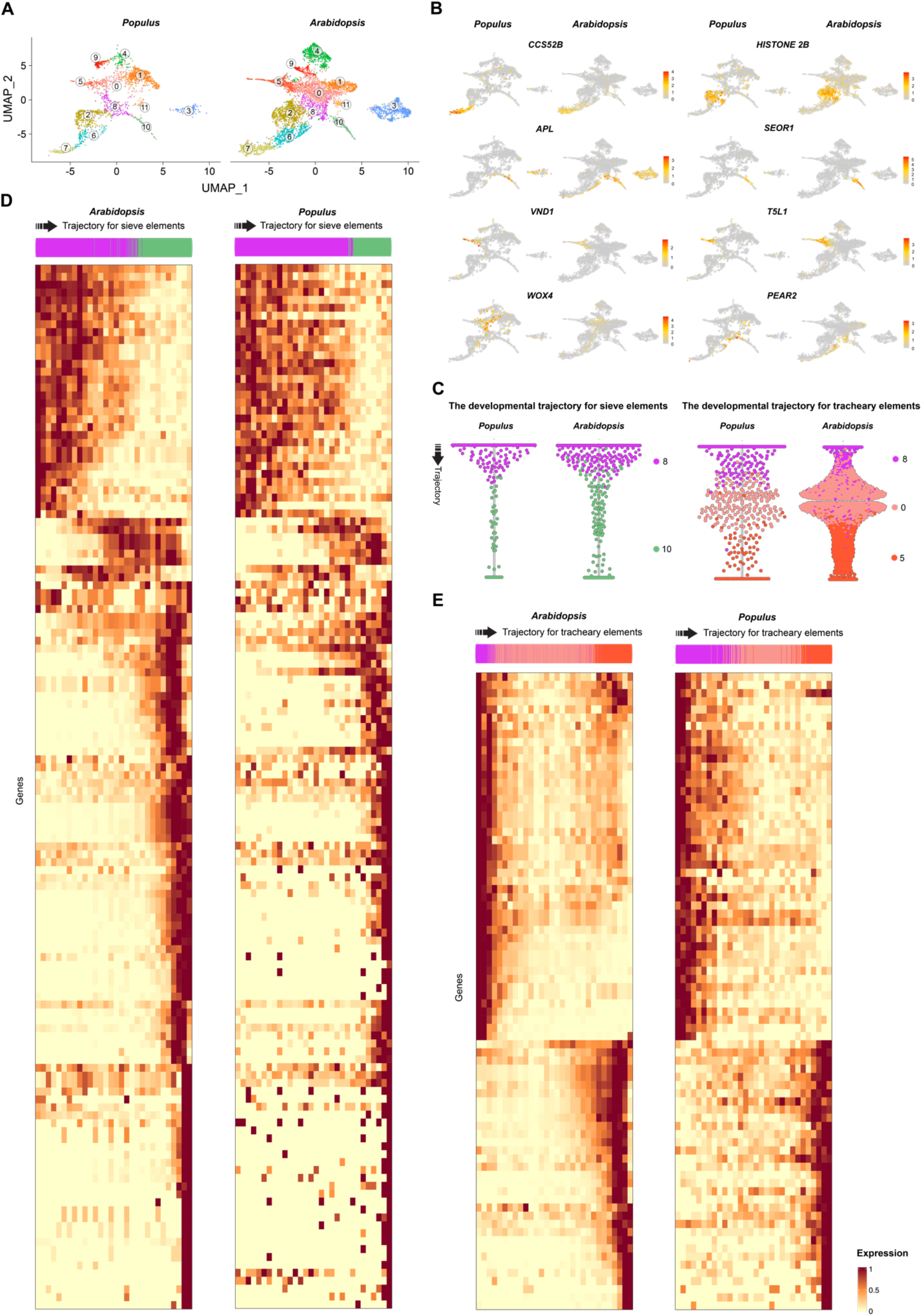
Characterization of the developmental trajectory of the primary vasculature development. **(A)** Clustering of the primary vasculature after the integration of *Arabidopsis* and *Populus* snRNA-seq data from the apex. **(B)** The expression of well-known markers identified the proliferating cells and primary phloem and tracheary elements. **(C)** Dynverse was used to run the trajectory for the primary phloem and tracheary elements in the *Populus*-*Arabidopsis* integrated data. Trade-seq was used to identify genes involved in the cell type specification. **(D)** Heatmap with the expression of the genes associated with the developmental trajectories of sieve elements in both *Populus* and *Arabidopsis* that shared the same expression pattern based on their positive correlated expression between species, calculated by dividing the trajectory into 30 bins. **(E)** Heatmap with the expression of the genes associated with the developmental trajectories of tracheary elements in both *Populus* and *Arabidopsis*, that shared the same expression pattern based on their positive correlated expression between species, calculated by dividing the trajectory into 30 bins.

### Comparison between *Populus* and *Arabidopsis* primary vasculature development at single-cell resolution

To uncover conserved and divergent regulatory mechanisms involved in the sieve and tracheary element differentiation, we used Dynverse to infer their developmental trajectories after the data integration. Then, we applied a generalized additive model regression implemented in tradeSeq (*28*) to rank genes according to the association between their expression and cell ordering along the trajectory. By extension, these genes are putative regulators of the cell differentiation process occurring during the formation of the sieve and tracheary elements, or genes controlled by these regulators. We identified 576 genes for the phloem and 191 genes for the xylem, associated with the developmental trajectory in *Populus* (FDR ≤ 0.01). For phloem, out of 576 genes, we could confidently identify (based on Table S6) the *Arabidopsis* homolog for 431 genes. Remarkedly, 60% of these genes (259 genes) were also significantly associated with the *Arabidopsis* trajectory (Table S7; Fig. S4). For xylem, 81% (129 genes) were also significantly associated in *Arabidopsis* (Table S8; Fig. S5). To identify which genes present the same expression pattern in both species during cell differentiation, we divided each phloem and tracheary element trajectories into 30 bins with an equal number of cells. Bins were constructed continuously along the trajectory such that the first bin contained the first ^1^/30 cells until the 30^th^ bin. Binning the cell data allowed us to quantify gene expression in regular segments along the trajectory in each species and, posteriorly, to calculate the gene expression correlation between *Populus* and *Arabidopsis* along the trajectories. Out of the 259 common genes for phloem development, 51% (132 genes) were significantly correlated (Corr. ≥ 0.5; FDR ≤ 0.01) between species (Table S9; Fig. 5C). Among 129 common genes for the xylem, 60% (78 genes) were correlated (Table S10; Fig. 5D). Correlated genes IDs and their expression along the trajectories are shown in Fig. S6. This comparative single-cell analysis highlighted the conservation of key regulatory modules that control sieve elements differentiation, previously identified in *Arabidopsis*, including *CLV3/EMBRYO SURROUNDING REGION 45* (*CLE45*) (*29*, *30*), *ALTERED PHLOEM DEVELOPMENT* (*APL*) (*31*), *SIEVE ELEMENT OCCLUSION-RELATED 1* (*SEOR1*), *HIGH CAMBIAL ACTIVITY2* (*HCA2*) (*32*, *33*), or *LATERAL ROOT DEVELOPMENT 3* (*LRD3*) (*34*) (Fig. S6, Table S9). Among highly correlated but less characterized conserved genes implicated in phloem differentiation, we detected *HOMEOBOX PROTEIN 33* (*HB33*), *NAC DOMAIN CONTAINING PROTEIN* 57 and 75 (*NAC057* and *NAC075*), or *LONESOME HIGHWAY LIKE 1* (*LHL1*).

Correlated genes for tracheary elements differentiation included well-known regulators of vasculature development such as *MONOPTEROS* (*MP*), *VASCULAR RELATED NAC-DOMAIN PROTEIN 1* (*VND1*), *ACAULIS 5* (*ACL5*), *PHLOEM INTERCALATED WITH XYLEM* (*PXY*), or *TARGET OF MONOTEROS 5-LIKE 1* (*T5L1*) (*1*, *21*, *24*).

The comparison between the *Arabidopsis* and *Populus* trajectories also resulted in the identification of genes exclusively associated with phloem and xylem differentiation in *Populus* (Tables S11 and S12; Figs. S7 and S8). We identified transcription factors related to phytohormones implicated in vascular formation, including auxin (*AUXIN RESPONSE FACTOR 2*) and cytokinin signaling (*CYTOKININ-RESPONSIVE GROWTH REGULATOR*), homeodomain transcription factors (*HOMEOBOX PROTEIN 16*), or NAC domain-containing transcription factors (*NAC089*) were also found only in the *Populus* trajectories.

Our results suggest that the transcriptional programs of the primary vasculature formation identified in the model species *Arabidopsis* are highly conserved in perennial woody plants. However, a subset of genes significantly related to vasculature differentiation appears to be unique to *Populus* in their association to this developmental program. This observation highlights our datasets’ relevance to identifying previously unexplored cell differentiation mechanisms in both model species. This pipeline can also be applied to compare other developmental programs occurring in the shoot apex, such as leaf and epidermis differentiation.

### Comparison between *Populus* and *Arabidopsis* procambium

The procambium remains one of the most understudied plant tissues due to the challenge of dissecting the cell population derived from cell division in the SAM but not yet differentiated into phloem and xylem. Here we separated the different vascular cell populations and used developmental trajectory analyses of the *Populus* and *Arabidopsis* integrated data to identify clusters 0 and 8 as those containing the precursors of phloem and xylem, the procambium cells (Fig. 4A). Cells in clusters 0 and 8 are transcriptionally similar to proliferating cells, but, unlike PC clusters, their marker genes are not involved in cell division. Eventually, these procambium cells divide to give rise to the primary xylem and phloem, or the vascular cambium in later stages of development of woody perennial species. The PC cluster identified in *Populus* likely contains procambial dividing cells ongoing their differentiation to phloem and xylem. However, in tune with the trajectory followed in *Arabidopsis*, (*1*), clusters 0 and 8 contain the steady procambial cells that are the precursors for the phloem and xylem formation. Hence, these two clusters are suitable for studying the biology of the vegetative shoot apex procambial cells.

**Figure 4.**
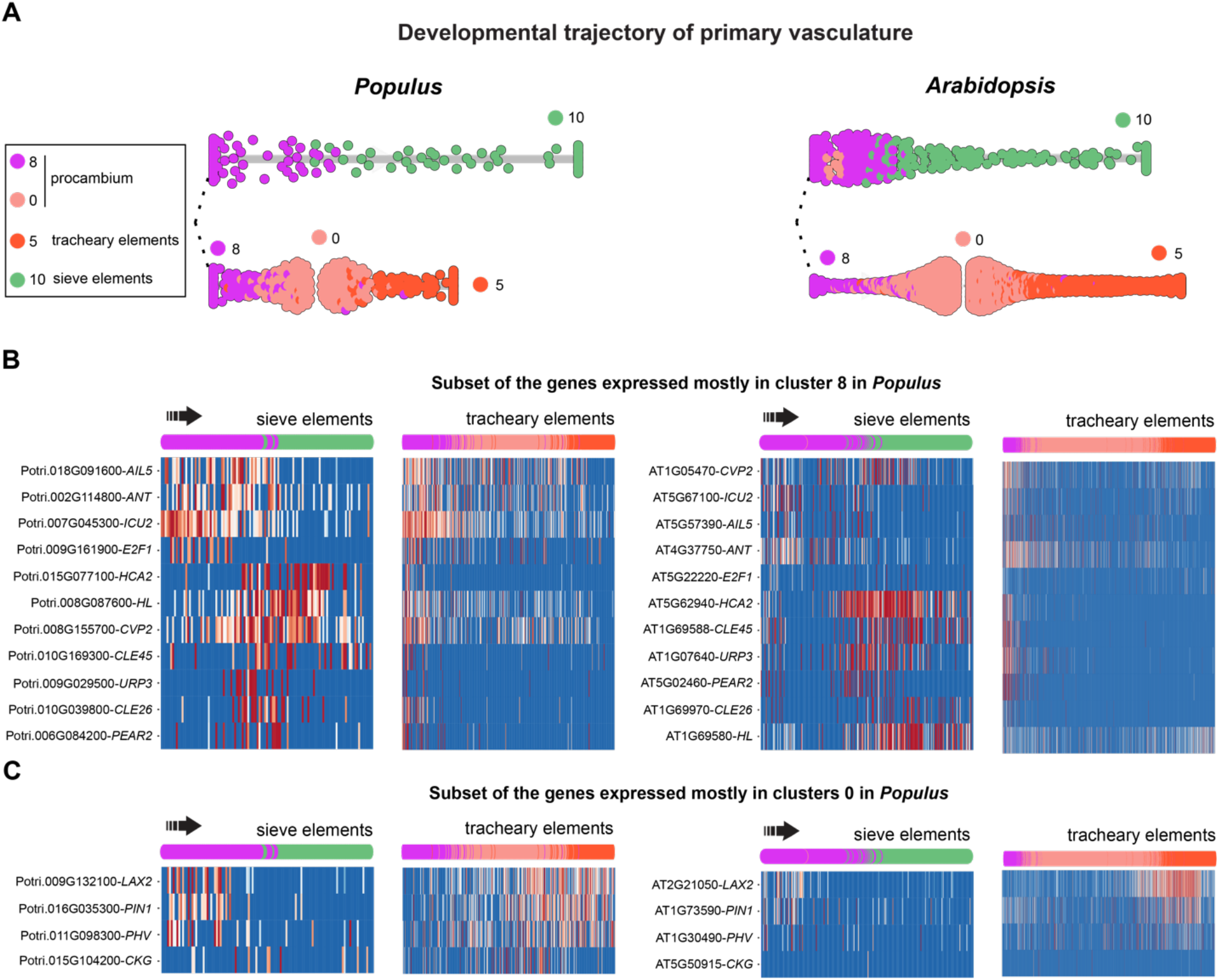
Identification of procambium cell regulatory genes. **(A)** Developmental trajectory of sieve (cluster10) and tracheary (cluster 5) elements from the procambial cells (cluster 0 and 8) inferred by dynberse. **(B)** A subset of the genes induced in cluster 8 of *Populus*, containing gene markers of vascular meristem such as *ANT*, and phloem cell initiation, such as *PEAR2*. Some of the patterns observed for *Populus* are shared in the *Arabidopsis* **(C)** Subset of genes induced in cluster 0 of *Populus*, containing genes involved in auxin signaling such as *LAX2* and *PIN1*.

To select genes potentially required for procambial cell maintenance, we identified differentially expressed genes (DEGs) in clusters 0 and 8 in *Populus* by comparing their transcript abundance to the other vascular cell clusters generated after the data integration (Fig. 3A, Fig. S3). We detected 417 DEGs, primarily expressed in these two clusters (FDR ≤ 0.05) (37 for cluster 0, 375 for cluster 8, and 5 that are common in both Table S13 and S14). For these 417 *Populus* genes, we identified 361 *Arabidopsis* homologs, of which 164 were also significantly induced in the *Arabidopsis* procambium (clusters 0 and 8 of the integrated data, Table S15).

Among the genes induced in cluster 8, we found *AINTEGUMENTA* (*ANT*), *AINTEGUMENTA-like 5* (*AIL5*), *INCURVATA 2* (*ICU2*) (Fig. 4B), and *FORKED 1* (*FDK1*) (Fig. 4B, Table S15). During secondary growth, *ANT* is mainly expressed in the vascular cambium in *Arabidopsis* and *Populus* stem (*35*–*37*). Other genes induced in cluster 8 showed a marked induction in the transition from procambium to phloem cells, as observed in *HIGH CAMBIAL ACTIVITY2* (*HCA2*), that promotes interfascicular cambium formation in *Arabidopsis* (*32*), *COTYLEDON VASCULAR PATTERN 2* (*CVP2*), which acts early in procambial patterning during embryogenesis in *Arabidopsis* (*38*), *PHLOEM EARLY DOF 2* (*PEAR2*), *CLE45*, or *UAS-TAGGED ROOT* PATTERNING3 (*URP3*). This evidence supports that cluster 8 contains procambial and phloem precursor cells (*39*, *40*). These expression patterns are shared with *Arabidopsis* (Fig. 4B, Table S15).

In agreement with the premise that cluster 0 contains procambial and xylem cells precursors in *Populus*, we identified *WOX4* and *PXY* expression significantly induced in this cluster. In the *Arabidopsis* root, auxin signaling is required for xylem differentiation (*41*). We found that poplar *PIN1* and *LAX2* are induced in cluster 0, and that their expression is expanded to the xylem cells (Fig. 4B), suggesting that these genes are involved in xylem differentiation. Remarkably, the expression of *MONOPTEROS* (*MP*), expressed in vascular cambium in the root (*41*), occurs later in the xylem development in the shoot apex of *Populus* and *Arabidopsis* (Fig. S6). This observation suggests a more specific role in xylem differentiation than procambium formation in the shoot apex primary vasculature. We also found a *Populus* homolog to *PHABULOSA* (*PHB*) induced in cluster 0, with a similar expression pattern to *PIN1*. *INCURVATA 4* (*ICU4/HB15*), which promotes vascular development in *Arabidopsis* (*42*), is expressed in procambial cells and is involved in xylem differentiation in *Zinnia elegans* (*43*) (Fig. 4B, Table S14). This gene is also induced in cluster 0, indicating that this cluster contains procambial cells and precursors of xylem development.

Our single-cell transcriptome analyses allowed us to infer the developmental trajectory of the primary vasculature in *Populus* and identify regulatory genes potentially involved in the xylem, phloem, and procambium development. We used the *Agrobacterium rhizogenes*-mediated poplar transformation root system and *promoter∷GUS* fusion to confirm that poplar *APL* (Potri.010G174100) is expressed early during vasculature differentiation in the root tip, specifically in the phloem (Fig. 5A). We also generated poplar stable transgenic lines to explore the *LAX2* (Potri.009G132100) promoter activity fused to GUS. GUS staining assay showed that, as predicted by the single-cell data, *LAX2* is expressed at the procambium and primary xylem during the early stages of vascular formation in the apex, when the vasculature is still present mainly in bundles (Fig. 5B). Further in the stem secondary growth, *LAX2* expression is restricted to the vascular cambium and the very first layers of the secondary xylem (Fig. 5B). These results highlight the accuracy of the cell clusters annotation and the vasculature developmental trajectories inferred for the *Populus* apex.

**Figure 5.**
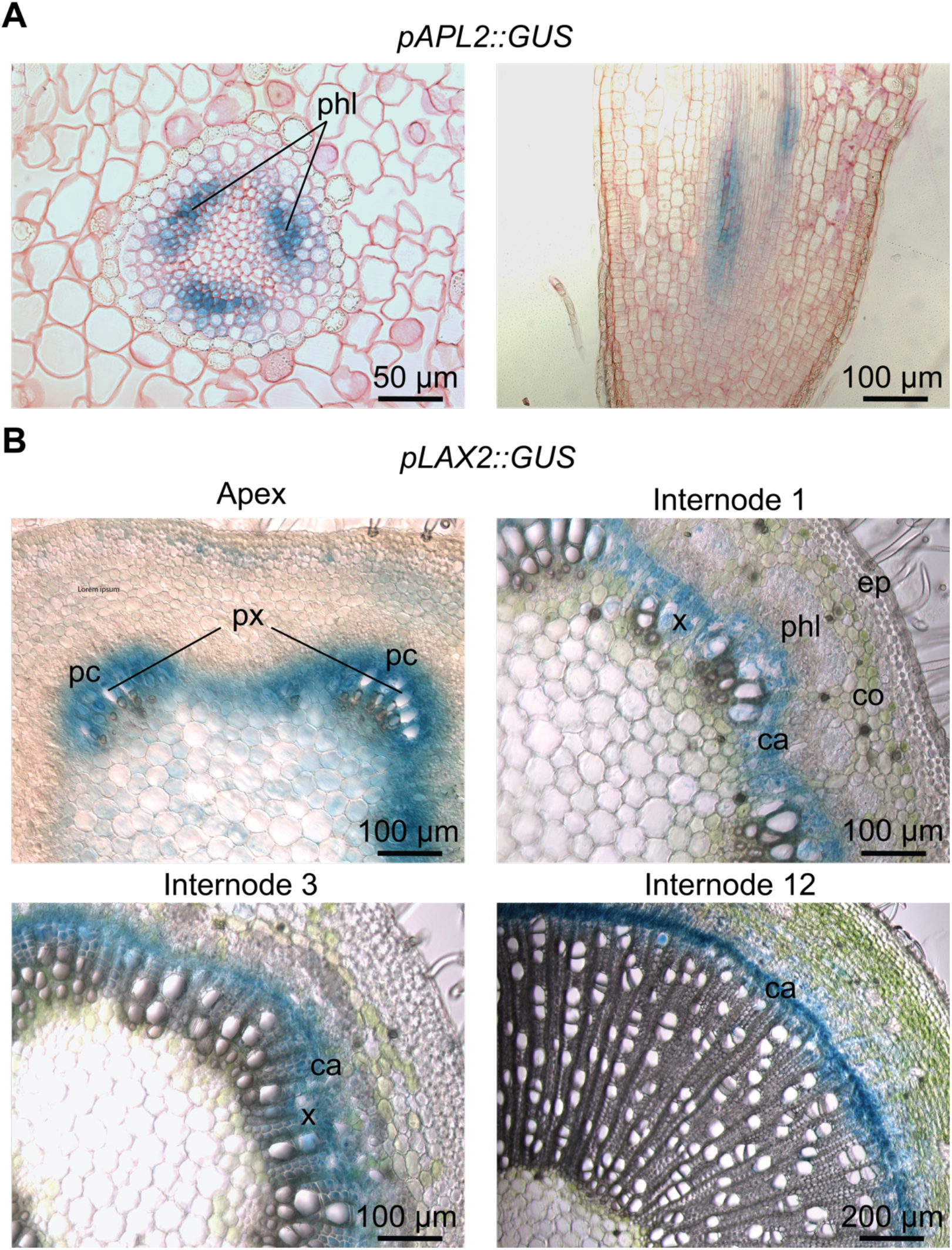
Validation of cell-type identity by *promoter∷GUS* fusions. **(A)** GUS activity under the *APL2* promoter, using the hairy root system, confirmed its expression very early during vasculature differentiation in the root tip, specifically in the phloem. **(B)** GUS activity under the *LAX2* promoter, in stable *Populus* transgenic lines, confirmed that *LAX2* is expressed at the procambium and primary xylem during the early stages of vascular formation when the vasculature is still present in bundles in the stem. During the stem secondary growth, *LAX2* expression is restricted to the vascular cambium and the very first layers of the secondary xylem. pc, procambium; px, primary xylem; phl, phloem; co, collenchyma; ep, epidermis; ca, vascular cambium; x, xylem.

## DISCUSSION

Single-cell RNA sequencing is revolutionizing the characterization of cell types and the developmental trajectories of cell lineages. Here we report the application of this technology to create a comprehensive single-cell gene expression atlas of the vegetative shoot apex of the woody perennial model species *Populus*.

Plant cells are surrounded by a cell wall, which differs in composition and thickness according to the species, tissue, developmental stage, and environmental conditions. The cell wall creates a hurdle for single-cell analysis of plant tissues, requiring its removal through enzymatic digestion and protoplast generation. However, certain plant cell types (e.g., trichomes) resist protoplasting (*1*). More importantly, the protoplasting procedure can lead to abrupt changes in transcriptomes. Challenges to the isolation of individual plant cells are further increased when working with highly lignified tissues, such as *Populus* stem. This study used single nucleus RNA sequencing (snRNA-seq) to overcome the above problems. We show that the isolation and analysis of individual nuclei result in the detection of diverse cell types and numbers of transcripts to enable their discrimination at a resolution comparable with whole cells in plants. Moreover, we demonstrated that this procedure can be applied to lignified *Populus* stems (*13*). The reported transcriptomes of shoot apex procambial cells, in combination with the feasibility of our nuclei isolation protocol to explore the single-cell transcriptomic in the stem, constitute an unpreceded opportunity to dissect the missing link between primary and secondary growth, including how cambium develops from procambium at the initiation of secondary growth.

By sequencing the transcriptome of individual cells, we identified a similar cell population structure to the one observed in the *Arabidopsis* shoot. We traced the developmental trajectories for mesophyll, epidermis, and vasculature in *Populus*. Motivated by the similar cell type structure found between both species, we developed an analysis pipeline to compare cellular developmental programs occurring during shoot development in annual and perennial model species. For this comparison, we initially focused on the primary vasculature development. Our *promoter∷GUS* staining results demonstrate the accuracy of unsupervised cell clustering and trajectory inference analysis in the delineation of cell differentiation toward the vasculature development in the *Populus* and *Arabidopsis* integrated data. In the future, this analytical framework can be applied to uncover the degree of conservation between cellular developmental programs that originate tissues such as the epidermis and mesophyll. This approach can also discover genes showing divergent expression patterns between species, which may explain their evolutionary differences. For instance, the same framework could be applied to compare cell fate determination and lateral organs and vascular tissue pattern formation in monocots and dicots or between gymnosperms and angiosperms.

Our results show the power of scRNA-seq technology to discover gene function in *Populus*. The precise and comprehensive gene expression pattern revealed by scRNA-seq may help to predict phenotypic changes that occur at specific tissues or cells, and guide us to choose the tissue that we should closely examine after the alteration of the expression of cell type specific genes, thereby markedly increasing the success rate of reverse genetics.

One limitation of scRNA-seq in plants is to match the messenger RNA profile of a cell with its position within a tissue or organ. One possible solution is spatial transcriptomics that allows the characterization of gene expression in barcoded regions of individual tissue sections. Although this technology has not been extensively applied in plant tissues (*44*), when combined with scRNA-seq or snRNA-seq, it has the potential to answer a wide range of biological questions concerning positional information in plants. Finally, the 10× Genomics Chromium system sensitivity is limited compared to other methods such as SMART-seq2 and Fluidigm C1 (*45*). As for *Arabidopsis*, when using the same technology, our scRNA-seq is insufficient to cluster previously identified cell types (i.e., CLV3 domain) within the SAM, based on the absence of the detection of *CLV3* gene expression.

In summary, we have generated a gene expression map of the vegetative shoot apex at single-cell resolution for the perennial model species *Populus*. The definition of cells in distinct layers and functional zones of the *Populus* shoot apex now offers researchers the opportunity to investigate, at unprecedented resolution, how stem cells differentiate into distinct cell types in perennial species and compare those mechanisms with the vast knowledge generated in the annual plants model species *Arabidopsis*, in a tissue-specific manner. Moreover, our procedure and results constitute a new opportunity to investigate the missing link between primary and secondary growth in perennials by comparing the specific transcriptomes of procambium and vascular cambium. Finally, our results demonstrate that we can identify procambial cells and their cell fate. The transition from procambial to differentiated vascular cells is mediated by spatial doses of positional signals that modulate the cell identity. The application of the snRNA-seq procedures and analyses described here will contribute in the future to identify the nature of those signals that so far remain poorly characterized.

## MATERIALS AND METHODS

### Plant material and growth conditions

Shoot tips and stem cuttings from *in vitro* grown hybrid poplar (*Populus tremula* × *alba* INRA clone 717 1B4) were used as explants for shoot multiplication. The explants were cut into small pieces (about 10–15 mm long) and placed aseptically on Murashige and Skoog media with vitamins (PhytoTech Labs, catalog number: M519) supplemented with 0.1 g/L myo-inositol (PhytoTech Labs, catalog number: I703), 0.25 g/L MES (PhytoTech Labs, catalog number: M825), 2% (w/v) sucrose (PhytoTech Labs, catalog number: S391) and solidified with 0.8% agar (PhytoTech Labs, catalog number: A296). The pH of the medium was adjusted to 5.8 with KOH before autoclaving. Explants were grown in a growth chamber under long-day conditions (16 hours light/8 hours dark), 22°C, 65% relative humidity, and 100 μmol m−2 s−1 photosynthetic photon flux. After three weeks in culture, a 5 mm-long portion of the shoot apex from 20 plants was excised, and leaves were removed, leaving only leaf primordia.

### Nuclei isolation from Populus apex for single nuclei RNA-seq

To perform nuclei isolation, we followed the previously developed protocol for poplar shoot apices (*13*).

### Single nuclei cDNA and library preparation

Twenty thousand nuclei were loaded to the 10X Genomics Chromium to generate the snRNA-seq library, following the 10× Genomics Chromium Single Cell v3.1 protocol. The cDNA reaction was prepared as described previously (*13*).

### Sequencing

Before full-scale library sequencing, a preliminary run was performed in an Illumina iSeq (~ 4M reads) to evaluate the parameters that are indicative of the quality of a scRNA-seq library, including the estimated number of cells captured and the fraction of reads in cells (i.e., percentage of ambient or “leaked” RNA). The sequencing was performed using a standard program: 28 bp (Cell barcode and UMI) for read 1, 90 bp (cDNA) for read 2, 10 bp for I7 Index, and 10 bp for I5 Index. The assessment of RNA leakage was derived from the profile of the relationship between UMI counts and cell barcodes detected (*13*). If the library was considered suitable for full-scale sequencing (reads in cells > 50% and at least 3,000 cells captured at low sequencing depth), we proceeded to sequence in an Illumina NovaSeq6000. We obtained 502M reads mapped to the genome, which resulted in a sequencing saturation of 78.9 % and an average of 59,475 reads per nucleus. Sequencing was performed at the Interdisciplinary Center for Biotechnology Research at the University of Florida (Gainesville, FL, USA).

### Cell clustering

Cell clustering was performed using Asc-Seurat *(14)*. The parameters used for cell clustering using all cells or for re-clustering specific cell types are described in Fig. S1.

### Identification of novel Populus cell-type-specific markers and DEG in the clusters

Cluster-specific genes were identified by Asc-Seurat using the Wilcoxon rank-sum test with the default parameters ((lfc ≥ 0.25, p-value adjusted ≤ 0.05, and expressed in at least 10 % of the cells of the cluster).

### Integration pipeline for Arabidopsis and Populus single-cell expression data

We integrated the *Populus* shoot apex vasculature data (Fig. 2B, Fig. S2A) with the single-cell gene expression data of the shoot apex vasculature of *Arabidopsis* (Fig. 6A of (*1*)). The one-to-one homology mapping of *P. trichocarpa* and *A. thaliana* genes was obtained from an extensive gene orthology generated from protein sequence data from 93 species genomes/transcriptomes. The input protein sequences included those from the *Medicago trucatula* v5 (*46*), *P. trichocarpa* v4.1 (*26*), and *A. thaliana* Araport 11 (*47*) genome assemblies, respectively. Gene ortho-groups and resolved (reconciled) gene trees were generated using the OrthoFinder 2 (*48*) framework, based on the blastp (*49*) (v2.2.28) and MCL (*50*) (v14-137) methodologies for gene orthology inference. The resolved gene trees from OrthoFinder 2 were further parsed by last common ancestor (LCA) duplication events with respect to the tree of 93 species to highlight better recent evolutionary dynamics across the species of interest in a refined set of parsed ortho-groups. For each parsed ortho-group (sub-tree) we checked if there was one gene member from both *P. trichocarpa* and *A. thaliana*, and identified such pairs of genes as one-to-one mappings. This analysis resulted in 9,827 high-confidence mappings for subsequent analysis of the data presented in this work. The data integration was performed in Seurat, using the expression of these 9,827 genes, in addition to the expression of 15 well-known tissue-specific markers showing a specific expression pattern in our vasculature single-cell data (Table S5).

### Validation of the predicted marker genes by GUS staining analysis

#### Cloning

For cloning the DNA constructs used in the present study, the Golden Gate system was used (*51*). A 1,715 bp and 1,840 bp segment of the promoters (until the ATG codon) of hybrid poplar *LAX2* and *APL* genes, respectively, were amplified by PCR using a Phusion High-Fidelity DNA Polymerase (New England BioLabs, Ipswich, MA, USA). The 4-nucleotide overhangs and the BsaI target sites at 5’ strands at both borders were added by including these sequences in the primers. Promoter sequences were cloned in frame with the beta-glucuronidase (GUS) CDS (pICH75111; MoClo Plant Parts Kit) and the 35S terminator (pICH41414; MoClo Plant Parts Kit) into the cloning vector pICH47811 (MoClo Toolkit) by using the level 1 reaction.

To characterize *pLAX2∷GUS* fusion activity, we generated stable transgenic lines. To identify the positive events of the transformation, we used the hygromycin resistance genes under the Atact2 promoter (pICH87644; MoClo Plant Parts Kit) cloned in the pICH47802 vector (MoClo Toolkit). Both transcriptional units containing the marker gene for the transformation and GUS, respectively, were cloned together into the final expression vector pAGM4673 (MoClo Toolkit) using the Golden Gate level 2 reaction. *Agrobacterium tumefaciens*-mediated transformation was performed using the strain GV3101 in the *Populus tremula* × *alba* 717-1B4 genotype, using the protocol described in (*52*).

To characterize the *pAPL∷GUS* fusion activity, we generated hairy roots. The fluorescent selection marker consisting of the CaMV 35S constitutive promoter driving the expression of the tdTomato fluorescence protein cloned into the pICH47802 vector (MoClo Toolkit) (*53*) was used to select transformed roots. Both transcriptional units containing the marker gene for the transformation and GUS, respectively, were cloned together into the final expression vector pAGM4673 (MoClo Toolkit) by using the level 2 reaction. For hairy roots generation, we followed a protocol described previously (*54*, *55*). Leaves of *Populus tremula* × *alba* 717-1B4 genotype in vitro plants were transformed with *Agrobacterium rhizogenes* strain MSU440.

#### GUS staining and preparation of cross sections

Three-week-old *in vitro* transgenic plants containing the *pLAX2∷GUS* construct were transferred to pots as described in (*54*). After 4 weeks of growth under a long-day regimen (16 hr light/8 hr darkness), at 22 °C, 65% relative humidity, and 100 μmol m−2 s−1 photosynthetic photon flux, a 5 mm-long portion of the apex, internode 1, 3 and 12, were transferred to the GUS staining solution (5 nM potassium ferrocyanide, 5 nM potassium ferricyanide, 0.1 M Sodium Phosphate Buffer, 1 mM Sodium EDTA, 1% Triton X and 0.3 % X-Gluc (previously dissolved in N,N-Dimethylformamide)).

To evaluate the activity of *pAPL∷GUS*, *in vitro* grown transgenic hairy roots after 4 weeks from the transformation, selected as the ones emitting the tdTomato fluorescence under an Olympus MVX10 fluorescence stereo microscope, were transferred to the GUS staining solution.

Apices, stems or roots were incubated for 2-4 hours in the solution. Next, plant material was fixed in 4% formaldehyde as previously described (*53*). Stained and fixed roots were then embedded using the Kulzer Technovit 7100 resin (Emgrid Australia) following the manufacturers’ instructions. Sections (8-10 μm thick) were obtained using a Leica RM2045 microtome and stained with 0.1 % Ruthenium Red (Sigma-Aldrich) in PBS for 5 minutes. Apices and stems were embedded in 6% (w/v) agarose and 40-60 μm thick cross sections were obtained in a Leica VT1000S vibratome. The images were obtained at Zeiss Axioplan 2 microscope attached to a QImaging Retiga EXi Fast 1394 camera.

## Funding

This work was supported by the Department of Energy Office of Science Biological and Environmental Research (Grant DE-SC0018247) to MK.

## Author contributions

DC and MK designed the research; DC, PMT, WJP, and MK designed the methodology; DC, PMT, HWS, and CD performed the experiments; DC, WJP, SAK, and KMB performed data processing and analysis; SR and MK supervised the data analyses. CD provided material and technical advice.; DC and MK wrote the paper. PMT, WJP, SAK, KMB, HWS, CD, and SR reviewed and edited the paper.

## Competing interests

None declared.

## Data and materials availability

Sequencing raw data and Cell Ranger outputs have been deposited in NCBI’s Gene Expression Omnibus and are accessible through GEO Series accession number GSE190649.

## Notes

### Competing Interest Statement

The authors have declared no competing interest.

## REFERENCES

1. T.-Q. Zhang, Y. Chen, J.-W. Wang, A single-cell analysis of the Arabidopsis vegetative shoot apex. Developmental Cell. 56, 1056–1074.e8 (2021).

2. A. L. Holt, J. M. van Haperen, E. P. Groot, T. Laux, Signaling in shoot and flower meristems of Arabidopsis thaliana. Current Opinion in Plant Biology. 17, 96–102 (2014).

3. J. R. Dinneny, P. N. Benfey, Plant Stem Cell Niches: Standing the Test of Time. Cell. 132, 553–557 (2008).

4. R. K. Yadav, M. Tavakkoli, M. Xie, T. Girke, G. V. Reddy, A high-resolution gene expression map of the Arabidopsis shoot meristem stem cell niche. Development. 141, 2735–2744 (2014).

5. C. Tian, Y. Wang, H. Yu, J. He, J. Wang, B. Shi, Q. Du, N. J. Provart, E. M. Meyerowitz, Y. Jiao, A gene expression map of shoot domains reveals regulatory mechanisms. Nat Commun. 10, 141 (2019).

6. K. D. Birnbaum, Power in Numbers: Single-Cell RNA-Seq Strategies to Dissect Complex Tissues. Annu. Rev. Genet. 52, 203–221 (2018).

7. V. Mironova, J. Xu, A single-cell view of tissue regeneration in plants. Current Opinion in Plant Biology. 52, 149–154 (2019).

8. C. Rich-Griffin, A. Stechemesser, J. Finch, E. Lucas, S. Ott, P. Schäfer, Single-Cell Transcriptomics: A High-Resolution Avenue for Plant Functional Genomics. Trends in Plant Science. 25, 186–197 (2020).

9. K. H. Ryu, L. Huang, H. M. Kang, J. Schiefelbein, Single-Cell RNA Sequencing Resolves Molecular Relationships Among Individual Plant Cells. Plant Physiol. 179, 1444–1456 (2019).

10. T. Denyer, X. Ma, S. Klesen, E. Scacchi, K. Nieselt, M. C. P. Timmermans, Spatiotemporal Developmental Trajectories in the Arabidopsis Root Revealed Using High-Throughput Single-Cell RNA Sequencing. Developmental Cell. 48, 840–852.e5 (2019).

11. C. N. Shulse, B. J. Cole, D. Ciobanu, J. Lin, Y. Yoshinaga, M. Gouran, G. M. Turco, Y. Zhu, R. C. O’Malley, S. M. Brady, D. E. Dickel, High-Throughput Single-Cell Transcriptome Profiling of Plant Cell Types. Cell Reports. 27, 2241–2247.e4 (2019).

12. T.-Q. Zhang, Z.-G. Xu, G.-D. Shang, J.-W. Wang, A Single-Cell RNA Sequencing Profiles the Developmental Landscape of Arabidopsis Root. Molecular Plant. 12, 648–660 (2019).

13. D. Conde, P. M. Triozzi, K. M. Balmant, A. L. Doty, M. Miranda, A. Boullosa, H. W. Schmidt, W. J. Pereira, C. Dervinis, M. Kirst, A robust method of nuclei isolation for single-cell RNA sequencing of solid tissues from the plant genus Populus. PLoS ONE. 16(2021), doi:10.1371/journal.pone.0251149.

14. W. Pereira, F. Almeida, K. Balmant, D. Rodriguez, P. Triozzi, H. Schmidt, C. Dervinis, G. Pappas, M. Kirst, “Asc-Seurat – Analytical single-cell Seurat-based web application” (preprint, Bioinformatics, 2021), doi:10.1101/2021.03.19.436196.

15. C. B. Lopez-Anido, A. Vatén, N. K. Smoot, N. Sharma, V. Guo, Y. Gong, M. X. Anleu Gil, K. Weimer, D. C. Bergmann, “Single-Cell Resolution of Lineage Trajectories in the Arabidopsis Stomatal Lineage and Developing Leaf” (preprint, Plant Biology, 2020), doi:10.1101/2020.09.08.288498.

16. J.-L. Gallois, C. Woodward, G. V. Reddy, R. Sablowski, Combined SHOOT MERISTEMLESS and WUSCHEL trigger ectopic organogenesis in *Arabidopsis*. Development. 129, 3207–3217 (2002).

17. B. Rutjens, D. Bao, E. van Eck-Stouten, M. Brand, S. Smeekens, M. Proveniers, Shoot apical meristem function in Arabidopsis requires the combined activities of three BEL1-like homeodomain proteins. The Plant Journal. 58, 641–654 (2009).

18. N. Ung, H. M. S. Smith, Regulation of shoot meristem integrity during Arabidopsis vegetative development. Plant Signaling & Behavior. 6, 1250–1252 (2011).

19. M. Menges, L. Hennig, W. Gruissem, J. A. H. Murray, Cell Cycle-regulated Gene Expression in Arabidopsis. Journal of Biological Chemistry. 277, 41987–42002 (2002).

20. C. Goldy, J.-A. Pedroza-Garcia, N. Breakfield, T. Cools, R. Vena, P. N. Benfey, L. De Veylder, J. Palatnik, R. E. Rodriguez, The *Arabidopsis* GRAS-type SCL28 transcription factor controls the mitotic cell cycle and division plane orientation. Proc Natl Acad Sci USA. 118, e2005256118 (2021).

21. D. Shi, V. Jouannet, J. Agustí, V. Kaul, V. Levitsky, P. Sanchez, V. V. Mironova, T. Greb, Tissue-specific transcriptome profiling of the Arabidopsis inflorescence stem reveals local cellular signatures. The Plant Cell. 33, 200–223 (2021).

22. S. Dinant, A. M. Clark, Y. Zhu, F. Vilaine, J.-C. Palauqui, C. Kusiak, G. A. Thompson, Diversity of the Superfamily of Phloem Lectins (Phloem Protein 2) in Angiosperms. Plant Physiology. 131, 114–128 (2003).

23. B. Péret, K. Swarup, A. Ferguson, M. Seth, Y. Yang, S. Dhondt, N. James, I. Casimiro, P. Perry, A. Syed, H. Yang, J. Reemmer, E. Venison, C. Howells, M. A. Perez-Amador, J. Yun, J. Alonso, G. T. S. Beemster, L. Laplaze, A. Murphy, M. J. Bennett, E. Nielsen, R. Swarup, *AUX/LAX* Genes Encode a Family of Auxin Influx Transporters That Perform Distinct Functions during *Arabidopsis* Development. Plant Cell. 24, 2874–2885 (2012).

24. V. Jouannet, K. Brackmann, T. Greb, (Pro)cambium formation and proliferation: two sides of the same coin? Current Opinion in Plant Biology. 23, 54–60 (2015).

25. K. Street, D. Risso, R. B. Fletcher, D. Das, J. Ngai, N. Yosef, E. Purdom, S. Dudoit, Slingshot: cell lineage and pseudotime inference for single-cell transcriptomics. BMC Genomics. 19, 477 (2018).

26. G. A. Tuskan, S. DiFazio, S. Jansson, J. Bohlmann, I. Grigoriev, U. Hellsten, N. Putnam, S. Ralph, S. Rombauts, A. Salamov, J. Schein, L. Sterck, A. Aerts, R. R. Bhalerao, R. P. Bhalerao, D. Blaudez, W. Boerjan, A. Brun, A. Brunner, V. Busov, M. Campbell, J. Carlson, M. Chalot, J. Chapman, G.-L. Chen, D. Cooper, P. M. Coutinho, J. Couturier, S. Covert, Q. Cronk, R. Cunningham, J. Davis, S. Degroeve, A. Dejardin, C. dePamphilis, J. Detter, B. Dirks, I. Dubchak, S. Duplessis, J. Ehlting, B. Ellis, K. Gendler, D. Goodstein, M. Gribskov, J. Grimwood, A. Groover, L. Gunter, B. Hamberger, B. Heinze, Y. Helariutta, B. Henrissat, D. Holligan, R. Holt, W. Huang, N. Islam-Faridi, S. Jones, M. Jones-Rhoades, R. Jorgensen, C. Joshi, J. Kangasjarvi, J. Karlsson, C. Kelleher, R. Kirkpatrick, M. Kirst, A. Kohler, U. Kalluri, F. Larimer, J. Leebens-Mack, J.-C. Leple, P. Locascio, Y. Lou, S. Lucas, F. Martin, B. Montanini, C. Napoli, D. R. Nelson, C. Nelson, K. Nieminen, O. Nilsson, V. Pereda, G. Peter, R. Philippe, G. Pilate, A. Poliakov, J. Razumovskaya, P. Richardson, C. Rinaldi, K. Ritland, P. Rouze, D. Ryaboy, J. Schmutz, J. Schrader, B. Segerman, H. Shin, A. Siddiqui, F. Sterky, A. Terry, C.-J. Tsai, E. Uberbacher, P. Unneberg, J. Vahala, K. Wall, S. Wessler, G. Yang, T. Yin, C. Douglas, M. Marra, G. Sandberg, Y. Van de Peer, D. Rokhsar, The Genome of Black Cottonwood, Populus trichocarpa (Torr. & Gray). Science. 313, 1596–1604 (2006).

27. D. M. Goodstein, S. Shu, R. Howson, R. Neupane, R. D. Hayes, J. Fazo, T. Mitros, W. Dirks, U. Hellsten, N. Putnam, D. S. Rokhsar, Phytozome: a comparative platform for green plant genomics. Nucleic Acids Research. 40, D1178–D1186 (2012).

28. K. Van den Berge, H. Roux de Bézieux, K. Street, W. Saelens, R. Cannoodt, Y. Saeys, S. Dudoit, L. Clement, Trajectory-based differential expression analysis for single-cell sequencing data. Nat Commun. 11, 1201 (2020).

29. S. Depuydt, A. Rodriguez-Villalon, L. Santuari, C. Wyser-Rmili, L. Ragni, C. S. Hardtke, Suppression of Arabidopsis protophloem differentiation and root meristem growth by CLE45 requires the receptor-like kinase BAM3. Proceedings of the National Academy of Sciences. 110, 7074–7079 (2013).

30. N. Shimizu, T. Ishida, M. Yamada, S. Shigenobu, R. Tabata, A. Kinoshita, K. Yamaguchi, M. Hasebe, K. Mitsumasu, S. Sawa, BAM 1 and RECEPTOR - LIKE PROTEIN KINASE 2 constitute a signaling pathway and modulate CLE peptide-triggered growth inhibition in *ARABIDOPSIS* root. New Phytol. 208, 1104–1113 (2015).

31. M. Bonke, S. Thitamadee, A. P. Mähönen, M.-T. Hauser, Y. Helariutta, APL regulates vascular tissue identity in Arabidopsis. Nature. 426, 181–186 (2003).

32. Y. Guo, G. Qin, H. Gu, L.-J. Qu, *Dof5.6/HCA2*, a Dof Transcription Factor Gene, Regulates Interfascicular Cambium Formation and Vascular Tissue Development in *Arabidopsis*. The Plant Cell. 21, 3518–3534 (2009).

33. Y. Kondo, A. M. Nurani, C. Saito, Y. Ichihashi, M. Saito, K. Yamazaki, N. Mitsuda, M. Ohme-Takagi, H. Fukuda, Vascular Cell Induction Culture System Using Arabidopsis Leaves (VISUAL) Reveals the Sequential Differentiation of Sieve Element-Like Cells. The Plant Cell. 28, 1250–1262 (2016).

34. P. Ingram, J. Dettmer, Y. Helariutta, J. E. Malamy, Arabidopsis Lateral Root Development 3 is essential for early phloem development and function, and hence for normal root system development: LRD3 is essential for root phloem function. The Plant Journal. 68, 455–467 (2011).

35. W. Dewitte, S. Scofield, A. A. Alcasabas, S. C. Maughan, M. Menges, N. Braun, C. Collins, J. Nieuwland, E. Prinsen, V. Sundaresan, J. A. H. Murray, Arabidopsis CYCD3 D-type cyclins link cell proliferation and endocycles and are rate-limiting for cytokinin responses. Proceedings of the National Academy of Sciences. 104, 14537–14542 (2007).

36. J. Schrader, J. Nilsson, E. Mellerowicz, A. Berglund, P. Nilsson, M. Hertzberg, G. Sandberg, A High-Resolution Transcript Profile across the Wood-Forming Meristem of Poplar Identifies Potential Regulators of Cambial Stem Cell Identity[W]. The Plant Cell. 16, 2278–2292 (2004).

37. R. S. Randall, S. Miyashima, T. Blomster, J. Zhang, A. Elo, A. Karlberg, J. Immanen, K. Nieminen, J.-Y. Lee, T. Kakimoto, K. Blajecka, C. W. Melnyk, A. Alcasabas, C. Forzani, M. Matsumoto-Kitano, A. P. Mähönen, R. Bhalerao, W. Dewitte, Y. Helariutta, J. A. H. Murray, *AINTEGUMENTA* and the D-type cyclin *CYCD3;1* regulate root secondary growth and respond to cytokinins. Biology Open. 4, 1229–1236 (2015).

38. F. M. Carland, T. Nelson, *COTYLEDON VASCULAR PATTERN2* –Mediated Inositol (1,4,5) Triphosphate Signal Transduction Is Essential for Closed Venation Patterns of Arabidopsis Foliar Organs. Plant Cell. 16, 1263–1275 (2004).

39. S. Miyashima, P. Roszak, I. Sevilem, K. Toyokura, B. Blob, J. Heo, N. Mellor, H. Help-Rinta-Rahko, S. Otero, W. Smet, M. Boekschoten, G. Hooiveld, K. Hashimoto, O. Smetana, R. Siligato, E.-S. Wallner, A. P. Mähönen, Y. Kondo, C. W. Melnyk, T. Greb, K. Nakajima, R. Sozzani, A. Bishopp, B. De Rybel, Y. Helariutta, Mobile PEAR transcription factors integrate positional cues to prime cambial growth. Nature. 565, 490–494 (2019).

40. T. Waki, S. Miyashima, M. Nakanishi, Y. Ikeda, T. Hashimoto, K. Nakajima, A GAL4-based targeted activation tagging system in *Arabidopsis thaliana*. Plant J. 73, 357–367 (2013).

41. O. Smetana, R. Mäkilä, M. Lyu, A. Amiryousefi, F. Sánchez Rodríguez, M.-F. Wu, A. Solé-Gil, M. Leal Gavarrón, R. Siligato, S. Miyashima, P. Roszak, T. Blomster, J. W. Reed, S. Broholm, A. P. Mähönen, High levels of auxin signalling define the stem-cell organizer of the vascular cambium. Nature. 565, 485–489 (2019).

42. I. Ochando, S. Gonzalez-Reig, J.-J. Ripoll, A. Vera, A. Martinez-Laborda, Alteration of the shoot radial pattern in Arabidopsis thaliana by a gain-of-function allele of the class III HD-Zip gene INCURVATA4. Int. J. Dev. Biol. 52, 953–961 (2008).

43. K. Ohashi-Ito, H. Fukuda, HD-Zip III Homeobox Genes that Include a Novel Member, ZeHB-13 (Zinnia)/ATHB-15 (Arabidopsis), are Involved in Procambium and Xylem Cell Differentiation. Plant and Cell Physiology. 44, 1350–1358 (2003).

44. S. Giacomello, F. Salmén, B. K. Terebieniec, S. Vickovic, J. F. Navarro, A. Alexeyenko, J. Reimegård, L. S. McKee, C. Mannapperuma, V. Bulone, P. L. Ståhl, J. F. Sundström, N. R. Street, J. Lundeberg, Spatially resolved transcriptome profiling in model plant species. Nature Plants. 3, 17061 (2017).

45. C. Ziegenhain, B. Vieth, S. Parekh, B. Reinius, A. Guillaumet-Adkins, M. Smets, H. Leonhardt, H. Heyn, I. Hellmann, W. Enard, Comparative Analysis of Single-Cell RNA Sequencing Methods. Molecular Cell. 65, 631–643.e4 (2017).

46. Y. Pecrix, S. E. Staton, E. Sallet, C. Lelandais-Brière, S. Moreau, S. Carrère, T. Blein, M.-F. Jardinaud, D. Latrasse, M. Zouine, M. Zahm, J. Kreplak, B. Mayjonade, C. Satgé, M. Perez, S. Cauet, W. Marande, C. Chantry-Darmon, C. Lopez-Roques, O. Bouchez, A. Bérard, F. Debellé, S. Muños, A. Bendahmane, H. Bergès, A. Niebel, J. Buitink, F. Frugier, M. Benhamed, M. Crespi, J. Gouzy, P. Gamas, Whole-genome landscape of Medicago truncatula symbiotic genes. Nature Plants. 4, 1017–1025 (2018).

47. C. Cheng, V. Krishnakumar, A. P. Chan, F. Thibaud-Nissen, S. Schobel, C. D. Town, Araport11: a complete reannotation of the *Arabidopsis thaliana* reference genome. Plant J. 89, 789–804 (2017).

48. D. M. Emms, S. Kelly, OrthoFinder: phylogenetic orthology inference for comparative genomics. Genome Biol. 20, 238 (2019).

49. S. McGinnis, T. L. Madden, BLAST: at the core of a powerful and diverse set of sequence analysis tools. Nucleic Acids Research. 32, W20–W25 (2004).

50. A. J. Enright, An efficient algorithm for large-scale detection of protein families. Nucleic Acids Research. 30, 1575–1584 (2002).

51. C. Engler, M. Youles, R. Gruetzner, T.-M. Ehnert, S. Werner, J. D. G. Jones, N. J. Patron, S. Marillonnet, A Golden Gate Modular Cloning Toolbox for Plants. ACS Synth. Biol. 3, 839–843 (2014).

52. F. Gallardo, J. Fu, F. R. Cantón, A. García-Gutiérrez, F. M. Cánovas, E. G. Kirby, Expression of a conifer glutamine synthetase gene in transgenic poplar. Planta. 210, 19–26 (1999).

53. P. M. Triozzi, H. W. Schmidt, C. Dervinis, M. Kirst, D. Conde, Simple, efficient and open-source CRISPR/Cas9 strategy for multi-site genome editing in *Populus tremula* × *alba*. Tree Physiology, tpab066 (2021).

54. C. L. Ribeiro, D. Conde, K. M. Balmant, C. Dervinis, M. G. Johnson, A. P. McGrath, P. Szewczyk, F. Unda, C. A. Finegan, H. W. Schmidt, B. Miles, D. R. Drost, E. Novaes, C. A. Gonzalez-Benecke, G. F. Peter, J. G. Burleigh, T. A. Martin, S. D. Mansfield, G. Chang, N. J. Wickett, M. Kirst, The uncharacterized gene *EVE* contributes to vessel element dimensions in *Populus*. Proc Natl Acad Sci USA. 117, 5059–5066 (2020).

55. K. Yoshida, D. Ma, C. P. Constabel, The MYB182 Protein Down-Regulates Proanthocyanidin and Anthocyanin Biosynthesis in Poplar by Repressing Both Structural and Regulatory Flavonoid Genes. Plant Physiol. 167, 693–710 (2015).

