## Supplemental material for "Single-cell analysis resolves the genetic program of primary vascular system formation in hybrid poplar"

### Supplementary material

#### Supplementary tables:

**Supplementary Table 1 -** Summary of snRNA sequencing and quality control overview

| **Poplar apex** | |
|---|---|
| Number of Nuclei | 8,324 |
| Total Number of genes Detected | 31,214 |
| Mean Genes per Nucleus | 2,477 |
| Median Genes per Nucleus | 2,308 |
| Mean UMIs per Nucleus | 3,618 |
| Median UMIs per Nucleus | 3,206 |
| Mean UMIs per Gene | 965 |
| Median UMIs per Gene | 408 |

**Supplementary Figures:**
**Supplementary figure legends:**
**Figure S1. (A)** Pipeline followed in Asc-Seurat to cluster the individual nuclei transcriptomes obtained from the hybrid poplar vegetative shoot apex. **(B)** The mesophyll, epidermis, and vascular cells were re-clustered together with their corresponding proliferating cells (PC), following the parameters shown in this figure.

**Figure S2. (A)** Visualization of the *Populus* and *Arabidopsis* vascular tissue cell populations with the proliferating vascular cells by UMAP. Dots, individual cells; color, cell clusters. **(B)** The expression of well-known markers identified proliferating cells, xylem, and companion cells.

**Figure S3. (A)** Pipeline followed in Asc-Seurat to cluster the cells after the *Populus-Arabidopsis* integration of the apex vasculature data. **(B)** the number of cells identified in each cluster for Populus and Arabidopsis after the data integration. **(C)** dynberse was used to run the overall trajectory for vasculature in the *Populus-Arabidopsis* integrated data to identify the clusters involved in the sieve and tracheary elements differentiation.

**Figure S4.** Heatmap with the expression of the genes associated with the developmental trajectories of the sieve elements, common in both *Populus* and *Arabidopsis* species, at single-cell level resolution along the trajectory.

**Figure S5.** Heatmap with the expression of the genes associated with the developmental trajectories of the tracheary elements, common in both *Populus* and *Arabidopsis* species, at single-cell level resolution along the trajectory.

**Figure S6.** Heatmap with the expression of the genes associated with the developmental trajectories of sieve and tracheary elements in *Populus* and *Arabidopsis*, with the same expression pattern in both species based on their positive correlated expression (Corr. $\geq 0.5$; FDR $\leq 0.01$), calculated by dividing the trajectory into 30 bins.

**Figure S7.** Heatmap with the expression of the genes associated with the sieve elements developmental trajectory only in *Populus*, at single-cell level resolution along the trajectory.

**Figure S8.** Heatmap with the expression of the genes associated with the tracheary elements developmental trajectories only in *Populus*, at single-cell level resolution along the trajectory.

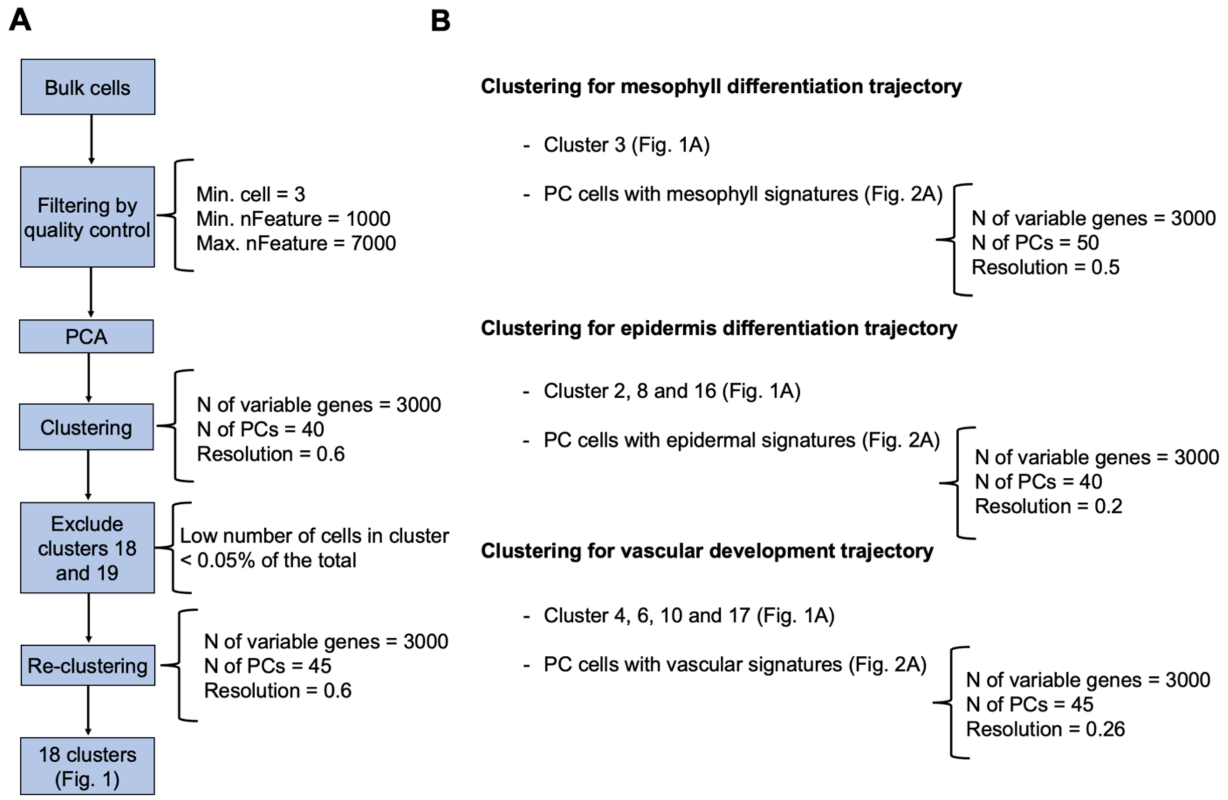
A
B
Bulk cells
Filtering by quality control
Min. cell = 3
Min. nFeature = 1000
Max. nFeature = 7000
PCA
Clustering
N of variable genes = 3000
N of PCs = 40
Resolution = 0.6
Exclude clusters 18 and 19
Low number of cells in cluster < 0.05% of the total
Re-clustering
N of variable genes = 3000
N of PCs = 45
Resolution = 0.6
18 clusters (Fig. 1)
Clustering for mesophyll differentiation trajectory
- Cluster 3 (Fig. 1A)
- PC cells with mesophyll signatures (Fig. 2A)
N of variable genes = 3000
N of PCs = 50
Resolution = 0.5
Clustering for epidermis differentiation trajectory
- Cluster 2, 8 and 16 (Fig. 1A)
- PC cells with epidermal signatures (Fig. 2A)
N of variable genes = 3000
N of PCs = 40
Resolution = 0.2
Clustering for vascular development trajectory
- Cluster 4, 6, 10 and 17 (Fig. 1A)
- PC cells with vascular signatures (Fig. 2A)
N of variable genes = 3000
N of PCs = 45
Resolution = 0.26

**Figure S1**

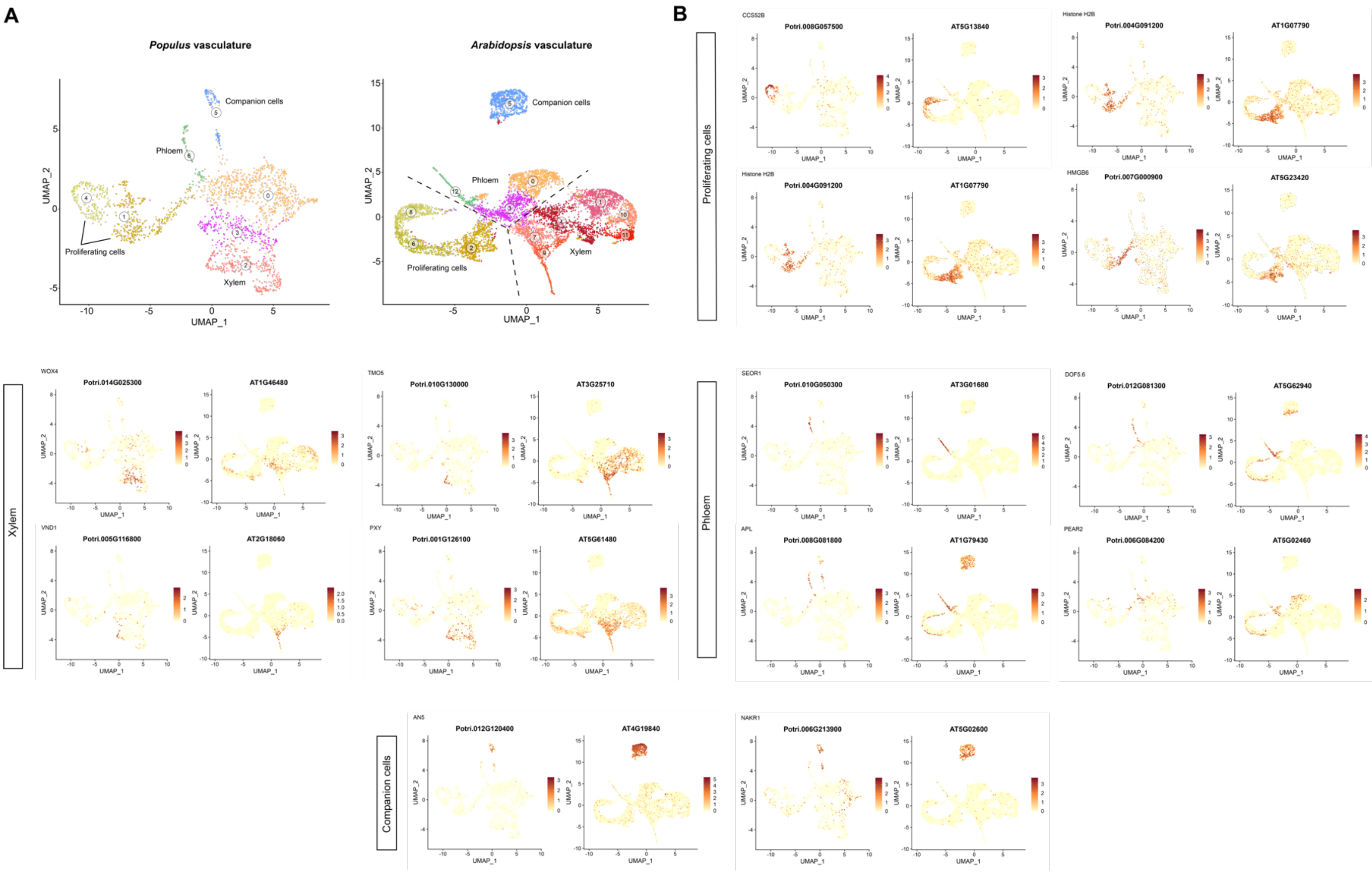
A
Populus vasculature
Arabidopsis vasculature
Companion cells
Phloem
Proliferating cells
Xylem
UMAP_1
UMAP_2
B
Proliferating cells
CCS52B
Potri.008G057500
AT5G13840
Histone H2B
Potri.004G091200
AT1G07790
HMGB6
Potri.007G000900
AT5G23420
Xylem
WOX4
Potri.014G025300
AT1G46480
TMO5
Potri.010G130000
AT3G25710
VND1
Potri.005G116800
AT2G18060
PXY
Potri.001G126100
AT5G61480
Phloem
SEOR1
Potri.010G050300
AT3G01680
DOF5.6
Potri.012G081300
AT5G62940
APL
Potri.008G081800
AT1G79430
PEAR2
Potri.006G084200
AT5G02460
Companion cells
AN5
Potri.012G120400
AT4G19840
NAKR1
Potri.006G213900
AT5G02600

**Figure S2**

**A**

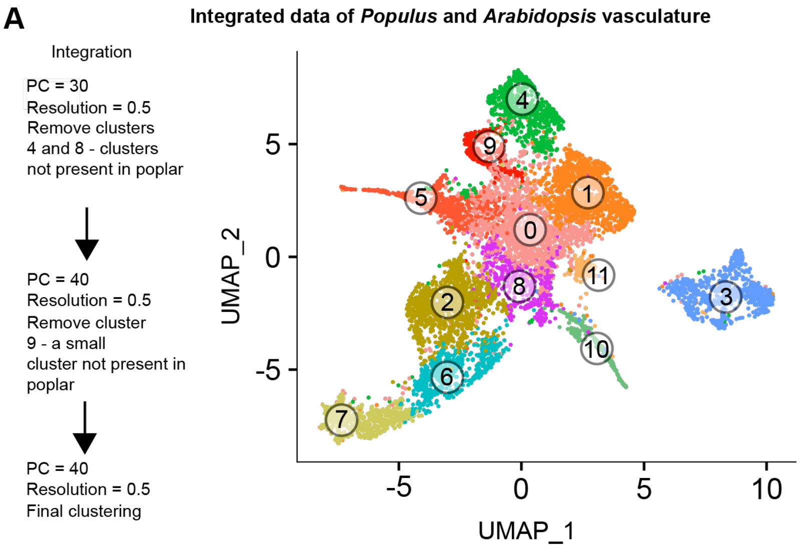


**B** Number of cells en each cluster

| Cluster | *Populus* | *Arabidopsis* |
|---|---|---|
| 0 | 195 | 1227 |
| 1 | 687 | 572 |
| 2 | 333 | 777 |
| 3 | 88 | 732 |
| 4 | 99 | 635 |
| 5 | 100 | 508 |
| 6 | 124 | 479 |
| 7 | 102 | 483 |
| 8 | 160 | 280 |
| 9 | 115 | 162 |
| 10 | 55 | 172 |
| 11 | 23 | 104 |

**C**

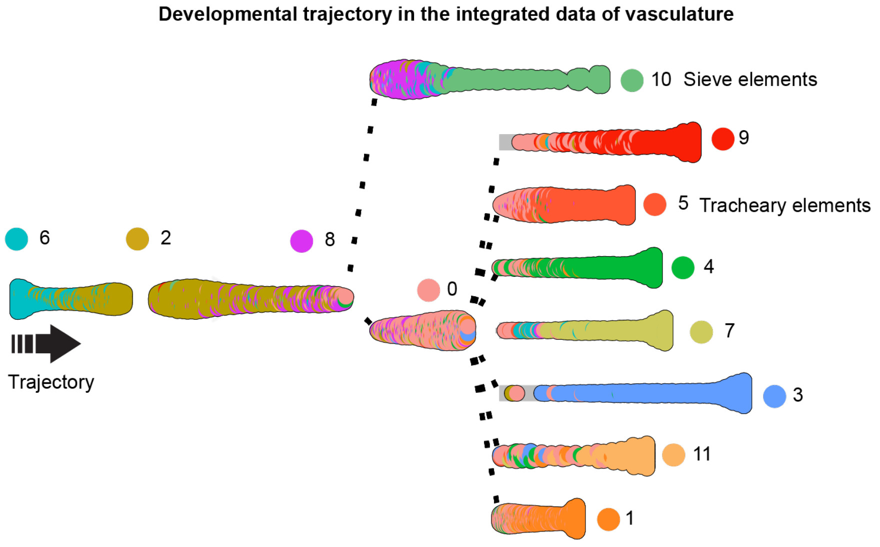


**Figure S3**

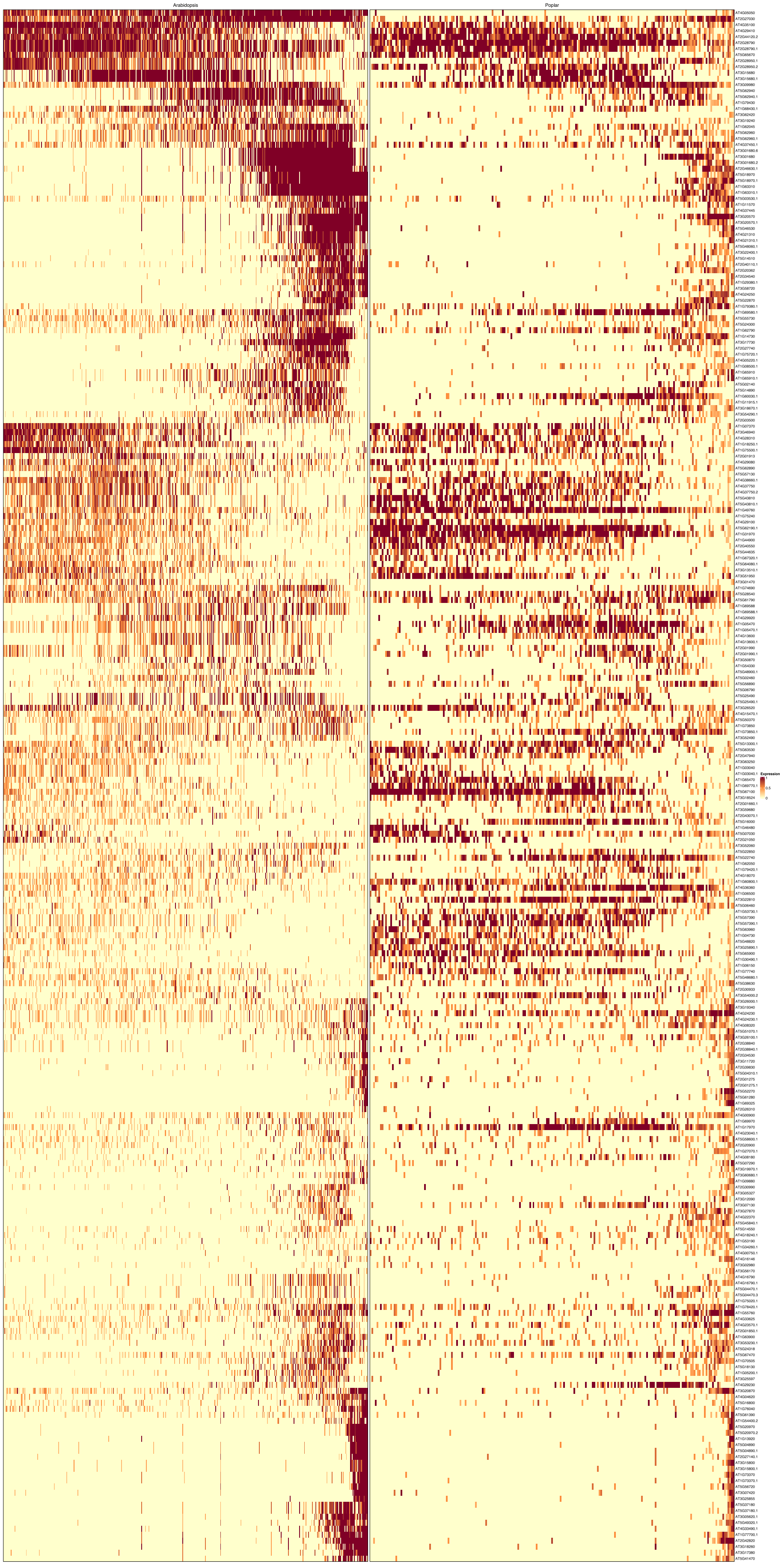


**Figure S4**

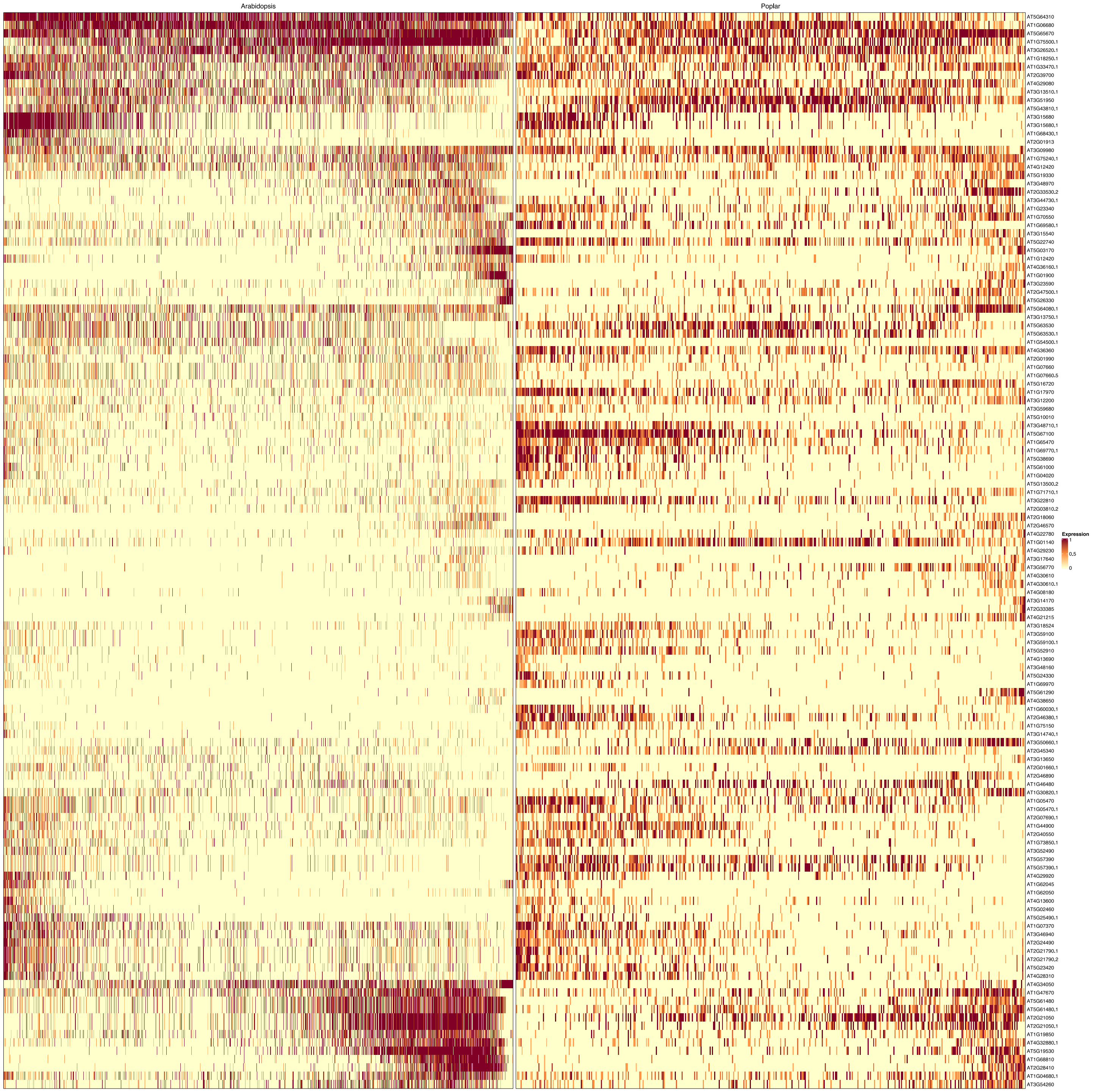
Arabidopsis
Poplar
AT5G64310
AT1G06680
AT5G65670
AT1G75500.1
AT3G26520.1
AT1G18250.1
AT1G33470.1
AT2G39700
AT4G29080
AT3G13510.1
AT3G51950
AT5G43810.1
AT3G15680
AT3G15680.1
AT1G68430.1
AT2G01913
AT3G09980
AT1G75240.1
AT4G12420
AT5G19330
AT3G48970
AT2G33530.2
AT3G44730.1
AT1G23340
AT1G70550
AT1G69580.1
AT3G15540
AT5G22740
AT5G03170
AT1G12420
AT4G36160.1
AT1G01900
AT3G23590
AT2G47500.1
AT5G26330
AT5G64080.1
AT3G13750.1
AT5G63530
AT5G63530.1
AT1G54500.1
AT4G36360
AT2G01990
AT1G07660
AT1G07660.5
AT5G16720
AT1G17970
AT3G12200
AT3G59680
AT5G10010
AT3G48710.1
AT5G67100
AT1G65470
AT1G69770.1
AT5G38690
AT5G61000
AT1G04020
AT5G13500.2
AT1G71710.1
AT3G22810
AT2G03810.2
AT2G18060
AT2G46570
AT4G22780
AT1G01140
AT4G29230
AT3G17640
AT3G56770
AT4G30610
AT4G30610.1
AT4G08180
AT3G14170
AT2G33385
AT4G21215
AT3G18524
AT3G59100
AT3G59100.1
AT5G52910
AT4G13690
AT3G48160
AT5G24330
AT1G69970
AT5G61290
AT4G38650
AT1G60030.1
AT2G46380.1
AT1G75150
AT3G14740.1
AT3G50660.1
AT2G45340
AT3G13650
AT2G01660.1
AT2G46890
AT1G46480
AT1G30820.1
AT1G05470
AT1G05470.1
AT2G07690.1
AT1G44900
AT2G40550
AT1G73850.1
AT3G52490
AT5G57390
AT5G57390.1
AT4G29920
AT1G62045
AT1G62050
AT4G13600
AT5G02460
AT5G25490.1
AT1G07370
AT3G46940
AT2G24490
AT2G21790.1
AT2G21790.2
AT5G23420
AT4G28310
AT4G34050
AT1G47670
AT5G61480
AT5G61480.1
AT2G21050
AT2G21050.1
AT1G19850
AT4G32880.1
AT5G19530
AT1G68810
AT2G28410
AT1G04680.1
AT3G54260
Expression
1
0.5
0

### Figure S5

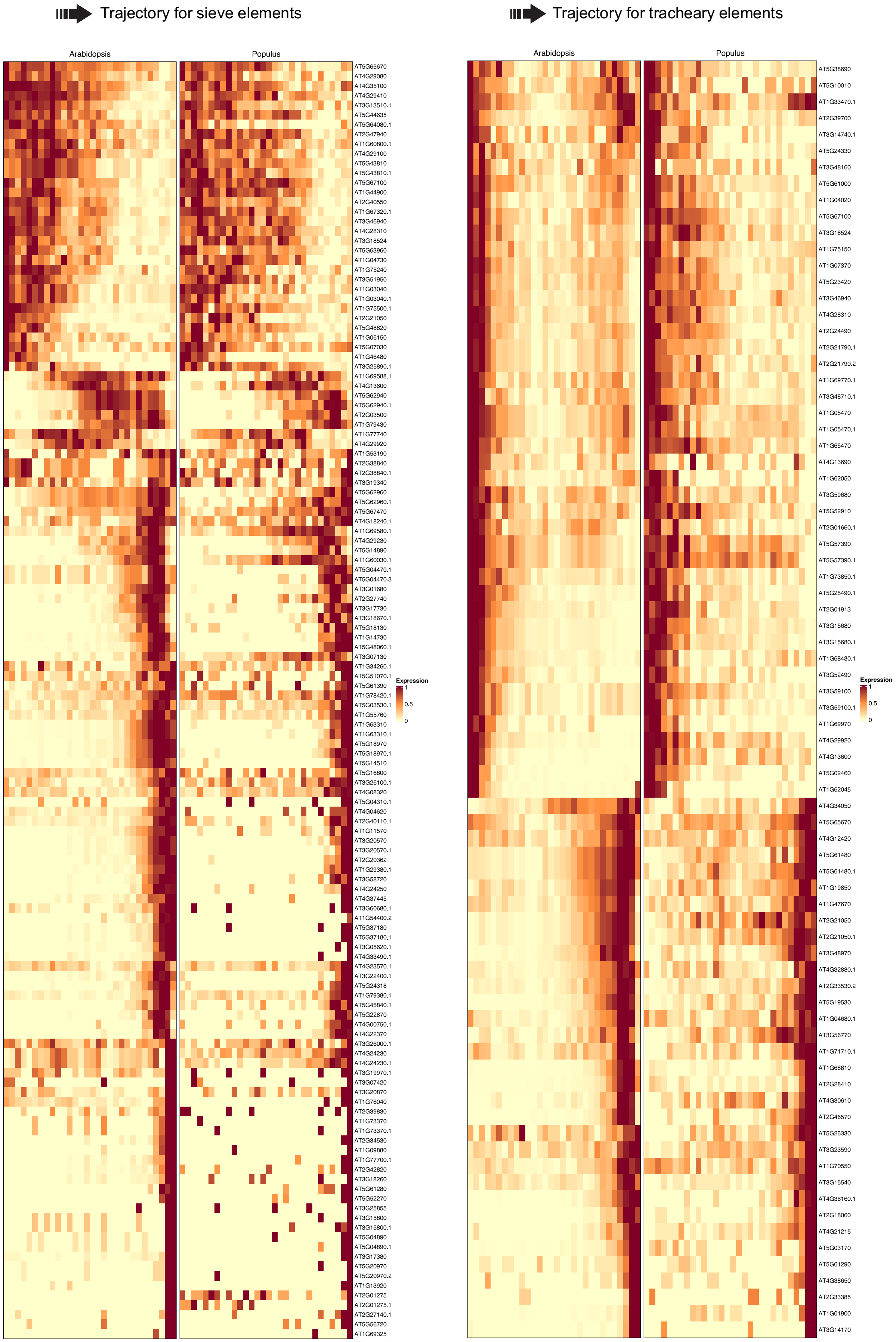
Trajectory for sieve elements
Arabidopsis
Populus
Expression
1
0.5
0
Trajectory for tracheary elements
Arabidopsis
Populus
Expression
1
0.5
0

**Figure S6**

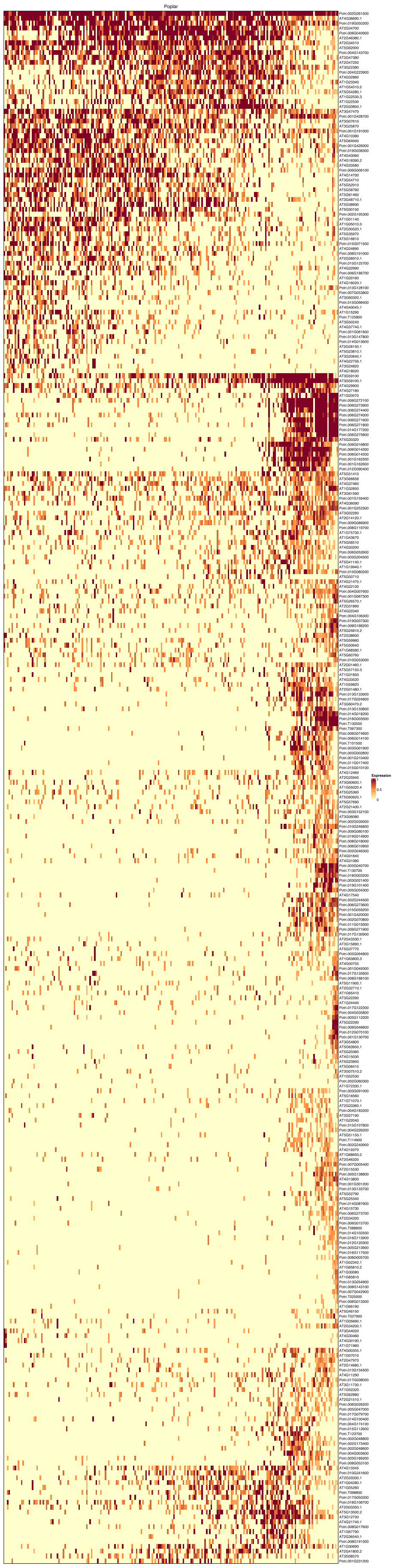


### Figure S7

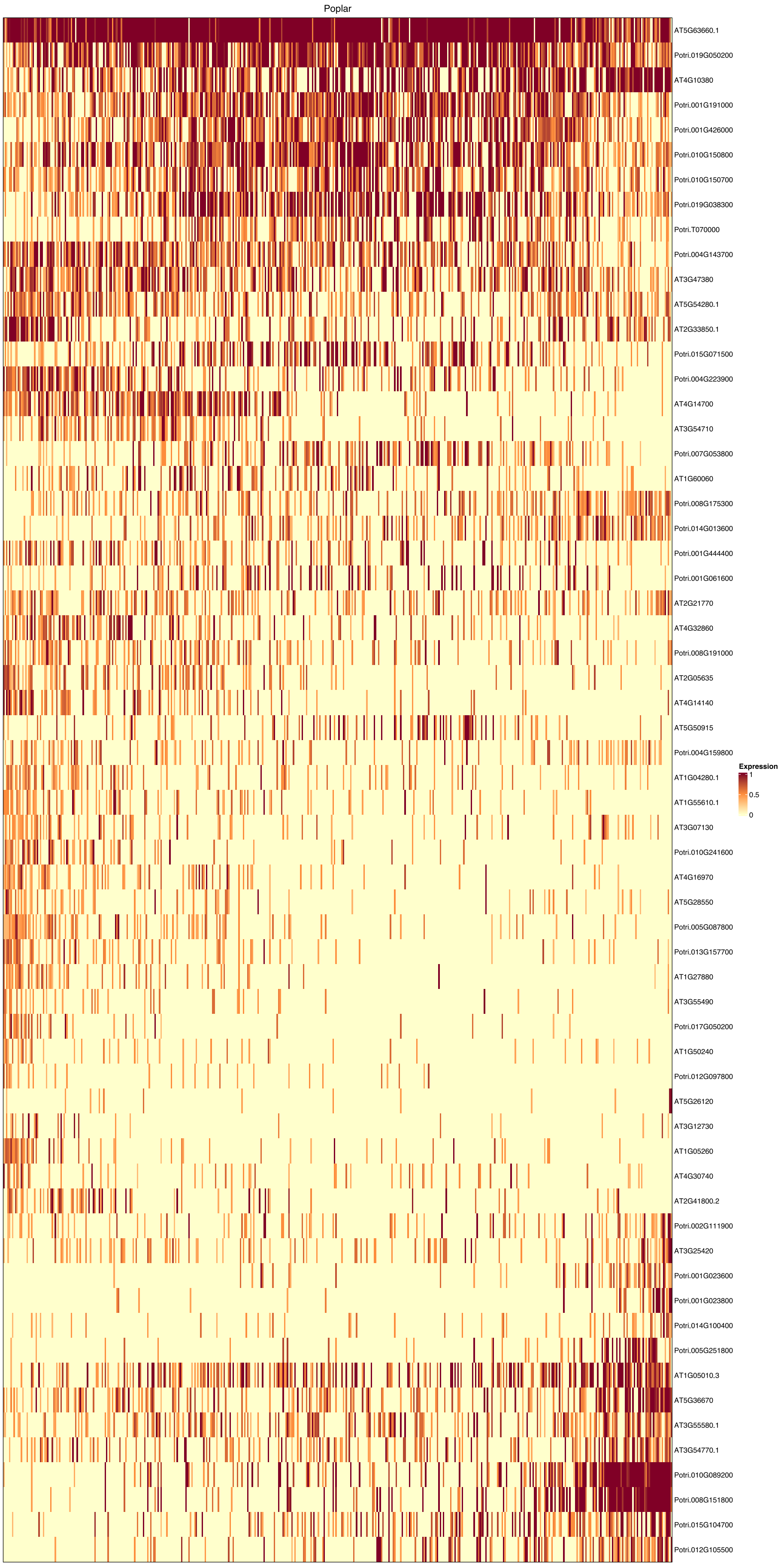
Poplar
AT5G63660.1
Potri.019G050200
AT4G10380
Potri.001G191000
Potri.001G426000
Potri.010G150800
Potri.010G150700
Potri.019G038300
Potri.T070000
Potri.004G143700
AT3G47380
AT5G54280.1
AT2G33850.1
Potri.015G071500
Potri.004G223900
AT4G14700
AT3G54710
Potri.007G053800
AT1G60060
Potri.008G175300
Potri.014G013600
Potri.001G444400
Potri.001G061600
AT2G21770
AT4G32860
Potri.008G191000
AT2G05635
AT4G14140
AT5G50915
Potri.004G159800
AT1G04280.1
AT1G55610.1
AT3G07130
Potri.010G241600
AT4G16970
AT5G28550
Potri.005G087800
Potri.013G157700
AT1G27880
AT3G55490
Potri.017G050200
AT1G50240
Potri.012G097800
AT5G26120
AT3G12730
AT1G05260
AT4G30740
AT2G41800.2
Potri.002G111900
AT3G25420
Potri.001G023600
Potri.001G023800
Potri.014G100400
Potri.005G251800
AT1G05010.3
AT5G36670
AT3G55580.1
AT3G54770.1
Potri.010G089200
Potri.008G151800
Potri.015G104700
Potri.012G105500
Expression
1
0.5
0

**Figure S8**
